# Dual Developmental Origins and Activity-dependent Specification of Mammalian Subplate Neurons

**DOI:** 10.64898/2026.06.12.731985

**Authors:** Takuma Kumamoto, Yuichiro Hara, Ryoka Katayama, Illia Aota, Hitomi Achiwa, Yurika Noguchi, Saki Gotoh-Saito, Ryoko Wada, Hiroyuki Hasegawa, Kazunori Nakajima, Hideya Kawaji, Chiaki Ohtaka-Maruyama

## Abstract

Subplate neurons (SpNs) are among the earliest-generated cortical neurons and are essential for neocortical circuit assembly. Despite this central role, they have long been considered a mammalian innovation, yet their evolutionary origin remains unresolved. Here, using comparative single-cell and spatial transcriptomics across amniotes (mice, chicks, and turtles), we identify two distinct developmental and evolutionary origins of SpNs: atypical SpNs (aSpNs), an Nr4a2-negative population conserved across amniotes and originating from the medial pallium, and mammalian-type SpNs (mSpNs), an Nr4a2-positive population preferentially expanded in mammals and arising from early-born cortical neurons. Cross-species analyses show that early-born pallial neurons in non-mammalian amniotes differentiate into thalamic input neurons, whereas this ancestral program is repurposed in mammals, with early-born neurons transiently adopting a subplate identity. We further show that this fate switch is controlled by Zbtb18 repression linked to thalamic input. Collectively, these findings establish a dual-origin model for SpNs and provide a unifying framework for understanding neocortical evolution.

**One-Sentence Summary:** Developmental rewiring of an ancestral input-neuron program gave rise to the mammalian subplate.

## Main

Subplate neurons (SpNs) are among the earliest-generated cortical neurons and play essential roles in establishing cortical circuits, including guiding thalamocortical projections and supporting early network activity (*1–6*). Disruption of SpNs impairs radial neuronal migration, cortical lamination, and the targeting of thalamocortical axons (TCAs) to layer 4, as well as ocular-dominance column formation, leading to long-lasting deficits in connectivity and function (*3, 6–9*).

Despite their vital role in early cortical circuit assembly, SpNs are widely regarded as a mammalian innovation (*1, 10, 11*). However, canonical SpN-associated markers, such as Connective Tissue Growth Factor (Ctgf/Ccn2) and Nuclear Receptor Subfamily 4 Group A Member 2 (Nr4a2), are also expressed in the sauropsid pallium (*12–14*), challenging the specificity of these genes as markers of a mammalian-specific cell type. Such ambiguity complicates the interpretation of molecular homology across species and how neocortical cell types evolved. This limitation is particularly critical because the emergence and diversification of cortical cell types are fundamentally linked to the expansion and increased complexity of mammalian cortical circuits. While earlier research has suggested that SpNs originate from multiple developmental sources, pointing to potential heterogeneity in their origins (*13*), a unified evolutionary and molecular framework remains elusive.

Together, these observations raise a key question: do SpNs represent a uniquely mammalian cell type, or do they reflect an evolutionarily conserved developmental program that has been differentially deployed across amniotes?

Here, we address this question using a comparative framework based on single-cell RNA sequencing (scRNA-seq) across the developing pallia of mammals (mice), birds (chicks), and reptiles (turtles) (Fig. 1A), complemented by spatial transcriptomic analyses (Visium, 10x Genomics). Our analysis reveals two distinct evolutionary roots of the SpN program and delineates their conserved and lineage-specific features.

**Fig. 1.**
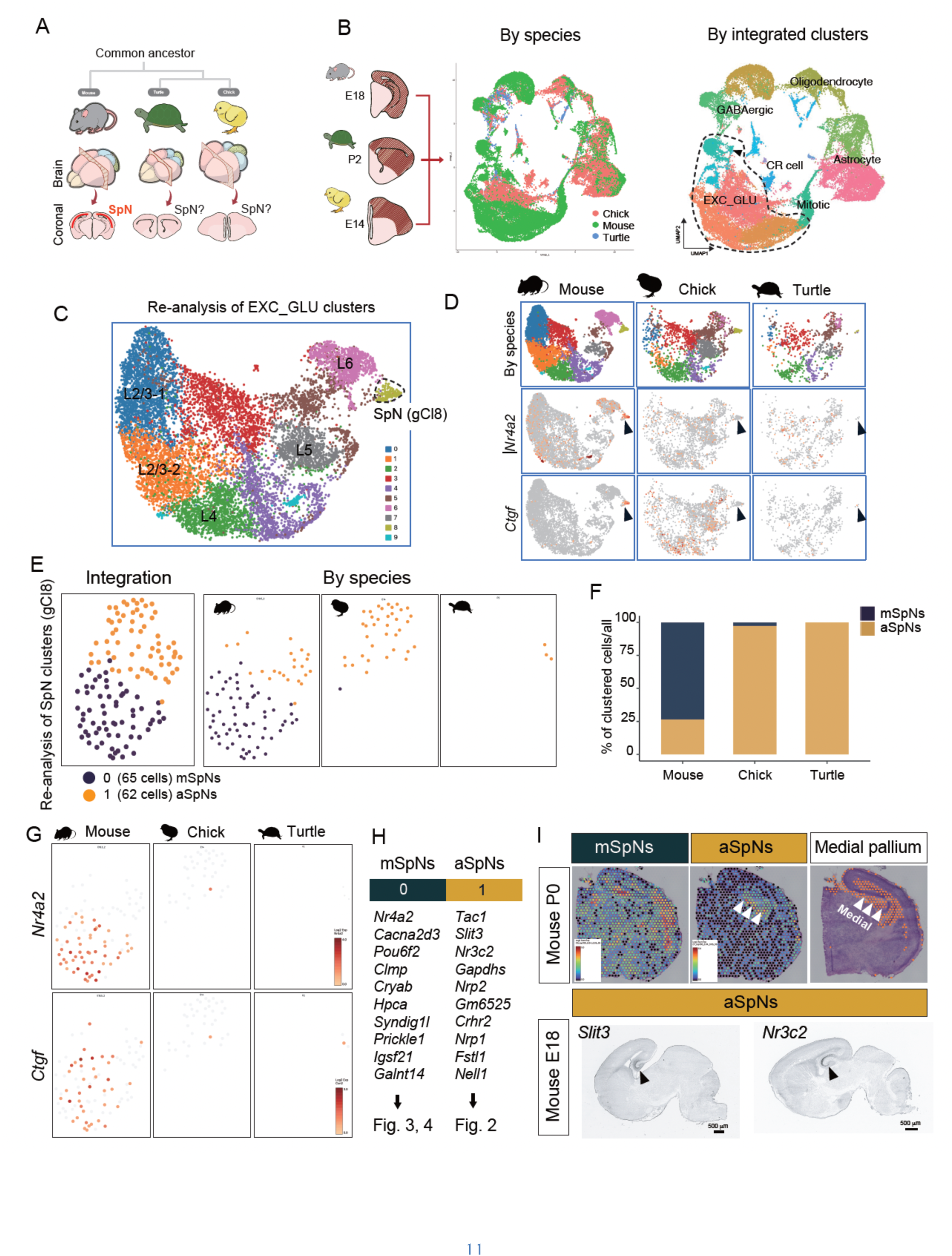
Two evolutionarily distinct subtypes of SpNs in amniote pallia. (**A**) Phylogenetic relationships among amniotes and their pallial brain structures. Red line shows SpNs in the mouse cortex. (**B**) Analyzing the flow of scRNA-seq and integration in this study. Uniform Manifold Approximation and Projection (UMAP) plot from the integrated dataset of the pallial regions, indicated by species (left) and clusters (right). (**C**) UMAP plot of the EXC_Glu clusters from (B). (**D**) Expression of the SpN markers, *Nr4a2* and *CTGF*, in the UMAP plot from (C), indicated by species. (**E**) UMAP plot of the SpN cluster (iCl8) from (D). Expression of the SpN markers, *Nr4a2* and *CTGF*, in the UMAP plot, indicated by species (right). Clusters 0 and 1 correspond to mSpNs and aSpNs, respectively. (**F**) The percentage of clustered cells from (E) in each species. (**G**) Expression of the SpN markers, *Nr4a2* and *CTGF*, in the UMAP plot from (E), indicated by species. (**H**) Top 10 genes listed in each cluster. (**I**) Spatial mapping of the average expression of the top 10 marker genes for clusters 0 and 1 onto the P0 mouse pallium using Visium. The medial pallial region is shown as a topographical reference. Bottom: *In situ* hybridization images from the E18 mouse brain, obtained from the Allen Brain Atlas, showing representative aSpN-enriched genes, Slit3 and Nr3c2. Arrowheads indicate enriched expression in the medial pallium.

### Two distinct SpN types emerge in the mammalian neocortex

To investigate the evolutionary origin of SpNs, we performed comparative scRNA-seq analyses across the pallia of mice, chicks, and turtles (Fig. S1). Cross-species integration—combining newly generated chick (E14) and turtle (P2) datasets with a published mouse (E18) reference using one-to-one orthologs—revealed a conserved glutamatergic population containing SpN-like cells in all species (cluster 9 (Cl9)), with a marked expansion in mice (Figs. 1B and S1–S3). Cell types were annotated based on established marker panels (e.g., Gad2 for GABAergic neurons, Neurod2 for glutamatergic neurons, Nr4a2 and Ctgf for SpN-associated populations, and lineage markers for glial cell types; see Methods).

Focusing on the glutamatergic compartment, re-clustering identified a distinct population corresponding to SpNs (glutamatergic cluster 8 (gCl8); Figs. 1C–E and S4). Subclustering of this population resulted in two distinct SpN subtypes (Fig. 1E). One subtype, detected primarily in mice, expressed canonical SpN markers, including *Nr4a2, Ctgf,* and *Cryab*. The other subtype, present across all three species, lacked *Nr4a2* expression and was enriched for genes such as *Tac1* and *Nr3c2* (Fig. 1F). We refer to these populations as mammalian-type SpNs (mSpNs) and atypical SpNs (aSpNs), corresponding to clusters 0 and 1, respectively (Fig. 1E,F). Feature plots of *Nr4a2* and *Ctgf* expression revealed that both genes are highly enriched in mSpNs, suggesting that they correspond to the canonical mammalian type (Fig. 1G). Differential expression analysis further defined these subtypes: mSpNs (Cl0) showed enrichment of SpN-associated genes, including *Nr4a2*, alongside additional canonical markers, such as *Cacna2d3* and *Cryab*, whereas aSpNs (Cl1) expressed several atypical genes, including *Tac1*, *Slit3*, and *Nr3c2* (Fig. 1H). These results distinguish a conserved aSpN population shared across amniotes from an mSpN population uniquely expanded in mice.

To examine the spatial distribution of these populations, we mapped subtype-specific gene signatures onto spatial transcriptomic datasets (Visium, 10x Genomics), including the chick E14 pallium and a previously published P0 mouse reference (*15*). This analysis revealed distinct localization patterns. The aSpN signature was restricted to the medial pallium in both mice and chicks (the term “medial” refers to its position within the pallium and does not imply hippocampal homology), whereas the mSpN signature localized to the neocortex in mice but was broadly distributed across the dorsal pallium in chicks (Figs. 1I and S7). These findings suggest distinct developmental origins for the two SpN subtypes and implicate the medial pallium as a source of aSpNs.

Consistent with this model, spatial transcriptomics in chicks showed that genes associated with the mouse subplate do not converge into a single spatial domain but are instead distributed across multiple pallial regions, indicating the absence of a discrete subplate-like structure (Figs. S6 and S7). To examine the evolutionary conservation of these programs, we queried pallial scRNA-seq data sampled from an amphibian *Pleurodeles waltl* (*16*). SpN-associated genes (*Ctgf*, *Nr4a2*, *Moxd1*) were broadly expressed across medial and dorsal pallial regions, whereas a subset of genes corresponding to the chick aSpN signature was enriched in the medial pallium. These results support a model in which medial pallium-derived SpNs represent an ancestral aSpN population, whereas dorsal pallium-derived SpNs correspond to a mammalian-specific mSpN program (Fig. S8).

### Medial pallium-derived aSpNs define an evolutionarily conserved program

To investigate whether aSpNs and mSpNs arise from distinct developmental origins in the mammalian neocortex, we performed lineage tracing by electroporating *^LiOn^CAG*∞*mScarlet-I3* and *CAG::GFP* (*17*) into the medial pallium at embryonic day (E)10 and traced labeled neurons to E15–E16. The medial pallium, including the Foxg1-negative cortical hem region (Fig. S9), gave rise to GFP+ neurons that migrated tangentially into the developing subplate and extended horizontally oriented processes (white box, Fig. 2A), consistent with the morphology of aSpNs. These neurons were MAP2-positive but lacked NR4A2 expression, in agreement with their molecular identity defined by scRNA-seq (Figs. 1G and 2B,C), and were derived predominantly from regions outside the Foxg1-negative cortical hem (Fig. S9). In contrast, SpNs derived from the cortical ventricular zone expressed NR4A2, corresponding to mSpNs (Fig. 2C).

**Fig. 2.**
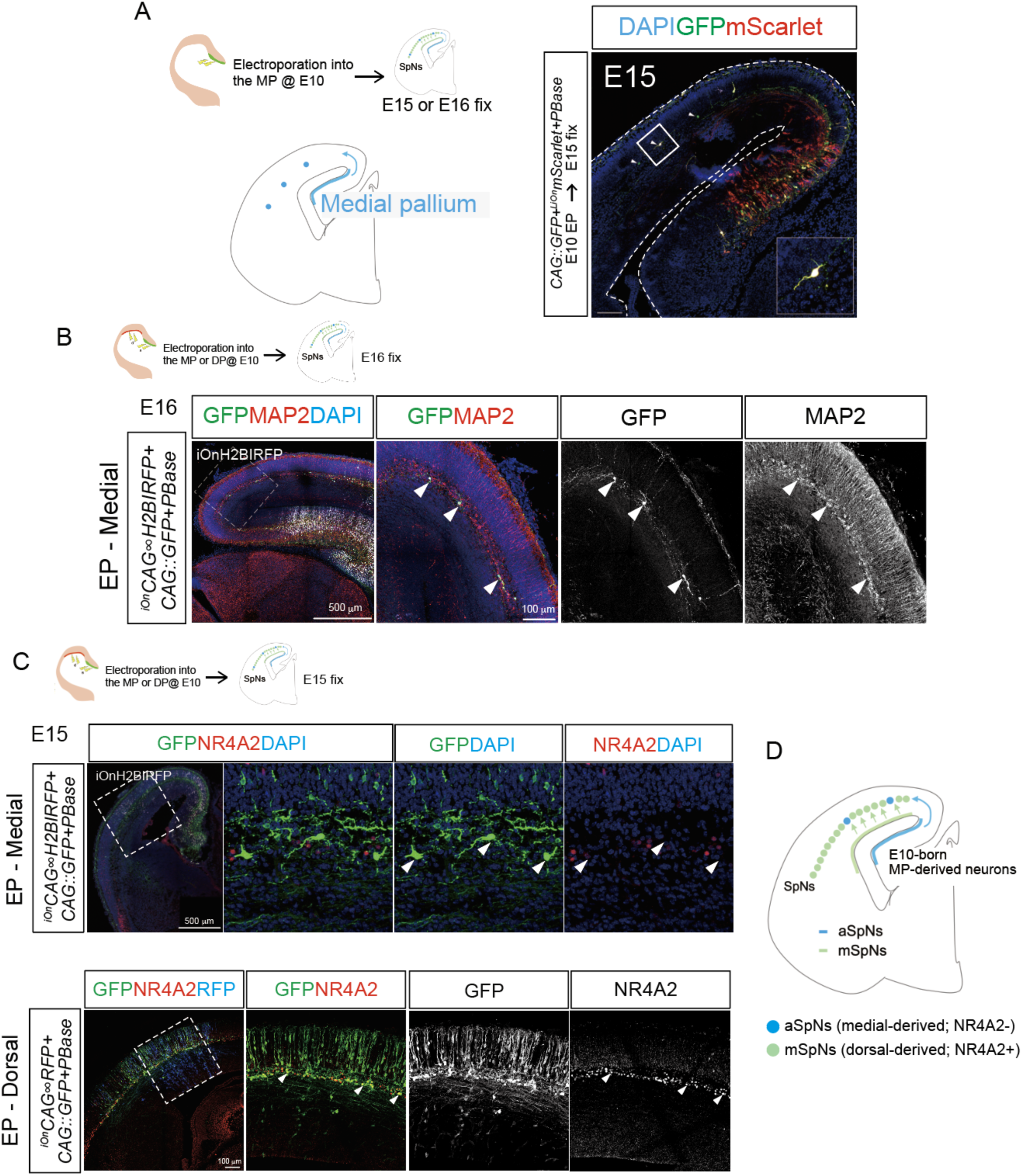
Origin of aSpNs in the mammalian neocortex. (**A**) Distribution of GFP-labeled neurons at E15 in the cortex. *CAG::GFP*, *^LiOn^CAG*∞*mScarlet-I3*, and *CAG::hyPBase* are electroporated at E10 in the mice medial pallium and fixed at E15. (**B**) SpN marker expression in GFP-labeled E10-born neurons. (Top) *CAG::GFP*, *^LiOn^CAG*∞*H2B-IRFP*, and *CAG::hyPBase* are co-electroporated at E10 in the mice medial cortex and fixed at E16. Green signal indicates GFP, red indicates MAP2, blue indicates DAPI, and white indicates IRFP. (**C**) (Top) *CAG::GFP*, *^LiOn^CAG*∞*H2B-IRFP*, and *CAG::hyPBase* are co-electroporated at E10 in the mice medial pallium and fixed at E15. Green signal indicates GFP, red indicates Nr4a2, blue indicates DAPI, and white indicates IRFP. (Bottom) *CAG::GFP*, *^iOn^CAG*∞*RFP*, and *CAG::hyPBase* are electroporated at E10 in the mice dorsal cortex and fixed at E15. Green signal indicates GFP, red indicates Nr4a2, and blue indicates RFP. (**D**) Summary of the spatial origin for SpNs in the mammalian neocortex.

These findings identify the medial pallium as a previously unrecognized source of SpNs in the mammalian neocortex. Specifically, it generates NR4A2-negative aSpNs that migrate tangentially into the subplate, complementing the NR4A2-positive mSpNs derived from the cortical ventricular zone and rostromedial telencephalic wall (*18*) (Fig. 2D).

### mSpN homologs align with an input-neuron program in non-mammalian amniotes

Next, we asked how mSpNs emerged in the mammalian neocortex. Because SpNs are among the earliest-born cortical neurons in mice (*19*) (Fig. S10), we examined early-born pallial neurons in non-mammalian amniotes as a shared developmental reference, using birth timing as a conserved axis for cross-species comparison (Fig. 3A). Previous studies, including ours, have mapped early-born chick neurons to outer pallial territories, particularly the entopallium and arcopallium (*20–22*) (pink, Fig. 3B). In the E14 chick Visium atlas, this region corresponded to a domain enriched for the SpN-associated gene Ctgf, motivating the use of Ctgf⁺ cells as candidate mSpN homologs (Fig. 3B). Ctgf⁺ neurons co-expressed the input-neuron marker Eag2/Kcnh5, indicating a clear input-neuron identity (Fig. 3D). *Ctgf* expression was already detectable at early developmental stages (from E11 in chicks; Fig. 3C), and birthdating analysis further showed that these neurons arise at the earliest stages of pallial neurogenesis (Fig. 3E).

**Fig. 3.**
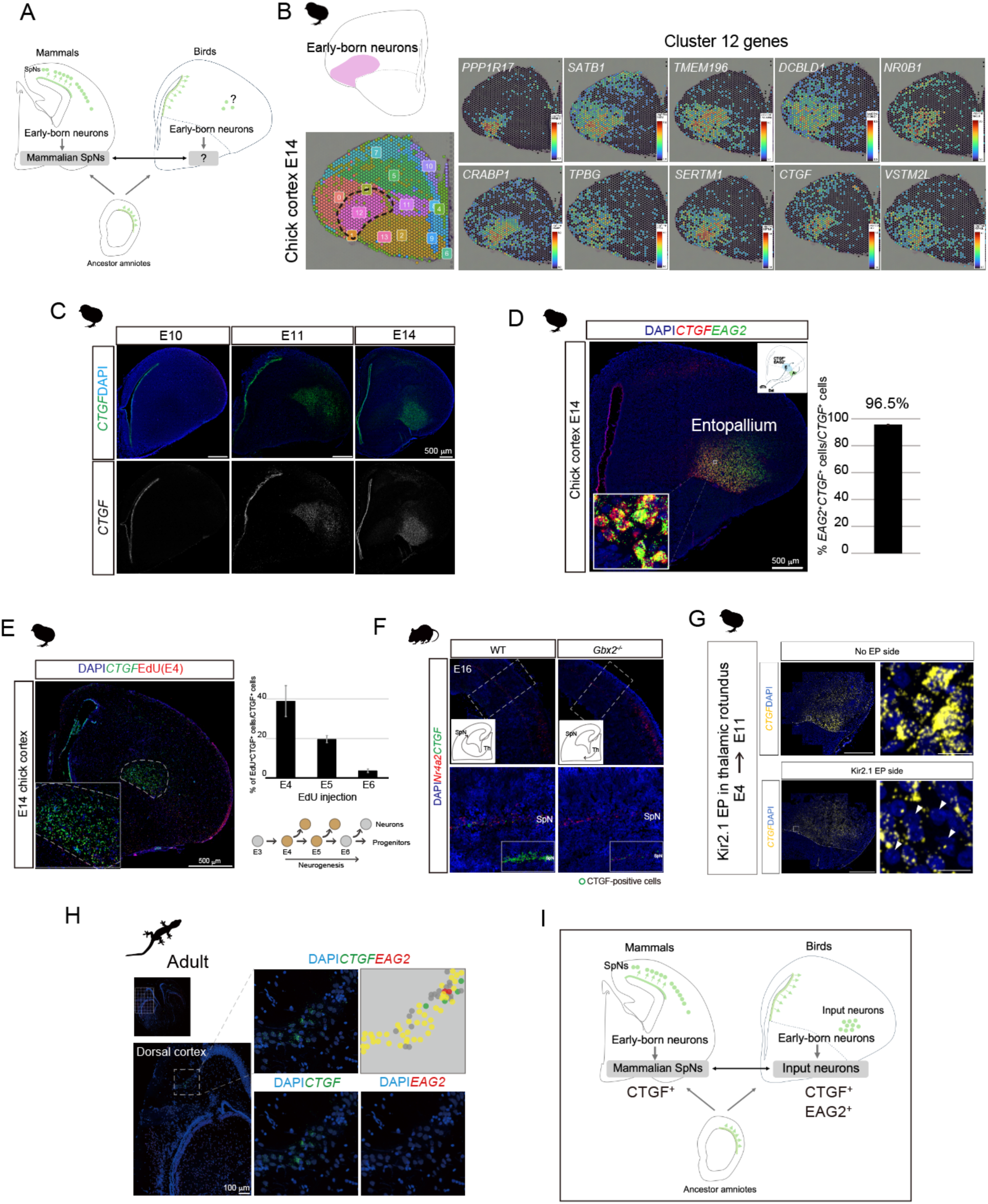
mSpNs are equivalent input neurons in non-mammalian pallia. (**A**) Scheme of SpN specification during cortical development in the mammalian neocortex. (**B**) (Top, left) Illustration of the early-born neuron distribution in the chick pallium, adapted from *Tsai et al. (1981), Fer (2024), and Katayama et al. (2024)*. (Bottom, left) Clustering of the chick E14 rostral Visium dataset. Black dotted line indicates an area where early-born neurons migrated. (Right) Top 10 significantly expressed genes in cluster 12 of the chick E14 rostral Visium data. **(C)** Temporal *Ctgf* expression pattern between E10 and E14 in the chick pallium. (**D**) Most *Ctgf*-positive cells in the chick entopallium are co-expressed with the input-neuron marker *Eag2*. (Right) Quantification of *Ctgf* and *Eag2* double-positive cells per all *Ctgf*-positive cells. (**E**) Birthdate analysis of *Ctgf*-positive entopallium neurons injected with EdU at E4, E5, and E6. (Middle) Magnified image of the white-dot region in the left image. (Right) Quantification of *Ctgf* and EdU double-positive cells per all *Ctgf*-positive cells in the entopallium. (**F**) TCA-derived input activity induces *Ctgf* expression in the mouse cortex. *Ctgf* and *Nr4a2* expression in the cortexes of wild-type (WT) (left) and *Gbx2* knockout (KO) (right) mice. Green indicates *Ctgf*, red indicates *Nr4a2*, and blue indicates DAPI. (**G**) TA-derived input activity induces *Ctgf* expression in the chick entopallium. *Ctgf* and *Nr4a2* expression in the no-EP side (top) and Kir2.1-EP side (bottom) in the chick entopallium. *CAG::Kir2.1*, *CAG::GFP*, and *^iOn^CAG*∞*RFP* are co-electroporated at E4 in the chick thalamic rotundus (Fig. S11). Yellow indicates CTGF, and blue indicates DAPI. (**H**) *Ctgf* and *Eag2* expression patterns in the adult gecko pallium. (**I**) Scheme of the summary in Fig. 3. Scale bars: (a, b) 500 μm. Nuclei are shown in blue using DAPI in all images.

Consistent with this identity, *Ctgf* expression localized to thalamorecipient regions such as the entopallium, intermediate hyperpallium apicale, and field L (Fig. S11), suggesting that neuronal activity induces Ctgf expression in the chick pallium.

Given that input-neuron specification in the chick entopallium depends on thalamic activity (*20*), we questioned whether *Ctgf* expression in mammalian SpNs is similarly regulated by thalamocortical input. In *Gbx2^-/-^* mice, which lack TCAs (*23*), cortical *Ctgf* expression was markedly reduced compared with that in wild-type controls (Fig. 3F), indicating regulation by thalamocortical input. This parallels the thalamic input-dependent specification observed in chicks (Figs. 3G and S12), supporting a shared regulatory logic across species. In addition, SpNs have been proposed to function as embryonic input neurons that receive thalamic input and exhibit early network activity (*2, 4*). Extending this analysis across species, *Ctgf*-positive neurons in reptiles also co-expressed Eag2, further supporting conservation of this input-neuron program across amniotes (Figs. 3H and S13). Collectively, these findings indicate that early-born pallial neurons across amniotes adopt a conserved input-neuron program, which is redeployed in mammals as mSpNs (Fig. 3I).

However, we must consider how this conserved input-neuron program is differentially deployed in mammals, where early-born neurons adopt a transient subplate identity while later-born neurons form the canonical thalamorecipient layer 4.

### Zbtb18 repression gates mSpN specification in the mammalian neocortex

To understand how early-born, thalamic input-receiving neurons adopt an SpN fate in mammals, we performed differential expression analyses across scRNA-seq datasets from chick entopallium cells (early-born input neurons), mouse layer 4 neurons (mammalian input neurons), and mouse SpNs (early-born mammalian neurons), thereby elucidating the regulators distinguishing these alternative fates. Specifically, the genes consistently downregulated in SpNs relative to input-neuron populations were identified to determine the factors underlying the divergence among these cell fates. ZBTB18 (RP58) emerged as a prime candidate, presenting a reciprocal expression pattern—expressed in both input-neuron populations (chick entopallium and mouse layer 4) but downregulated in SpNs (Fig. 4A). Notably, ZBTB18 encodes a transcription factor with established roles in cortical development, underscoring it as a candidate regulator underlying the divergence between input-neuron and SpN identities.

**Fig. 4.**
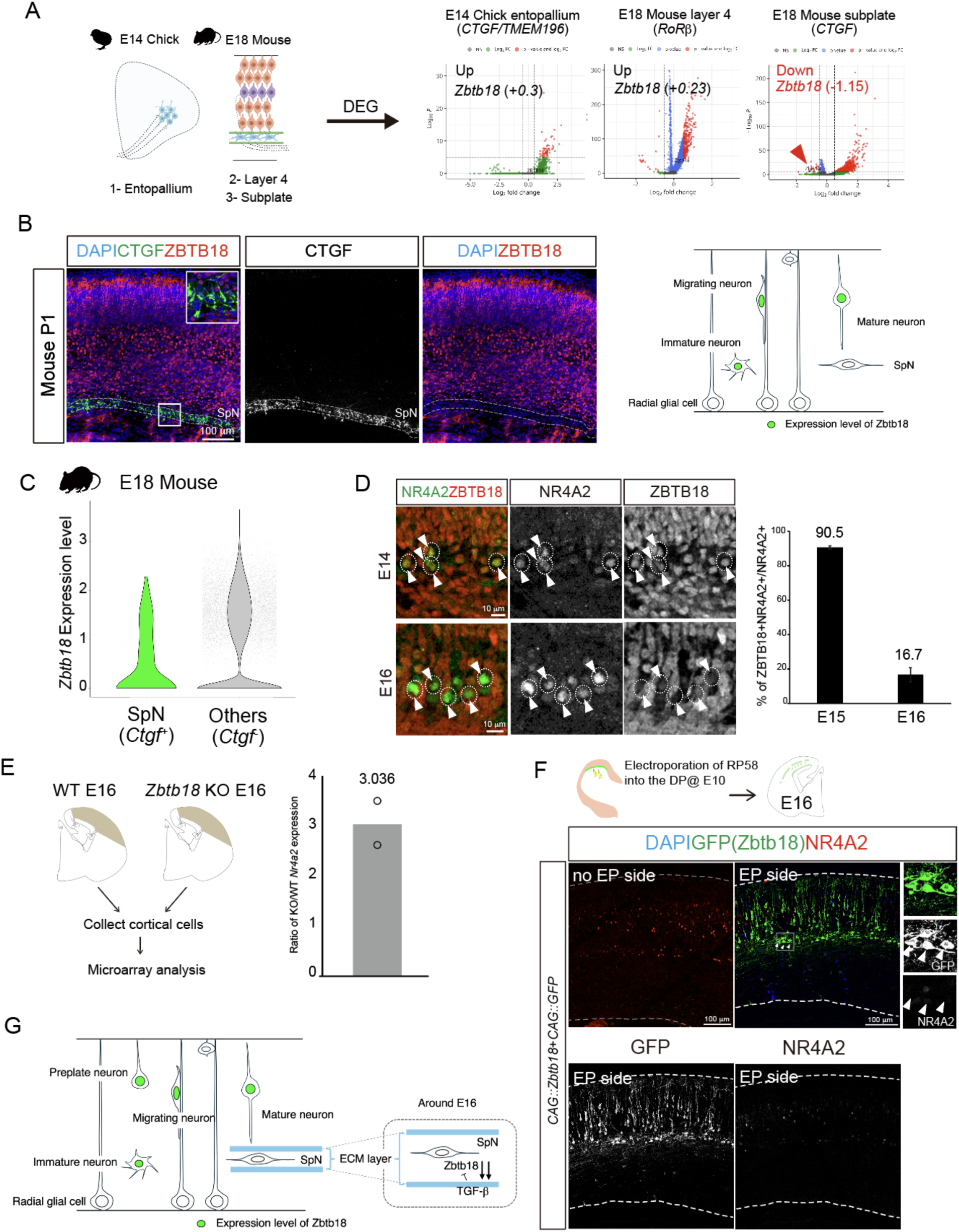
Zbtb18 repression contributes to mSpN specification in the mouse cortex. (**A**) Comparative transcriptomic analysis of the E14 chick entopallium and E18 mouse cortical neurons (layer 4 and SpNs). Volcano plots show differentially expressed genes for each comparison, with Zbtb18 indicated. (**B**) (Left) Immunostaining of CTGF and ZBTB18 in the P1 mouse cortex. (Right) Schematic of the Zbtb18 expression level during neural migration. (**C**) *Zbtb18* mRNA expression levels in SpNs compared with other cortical regions by scRNA-seq. **(D)** (Left) Immunostaining of NR4A2 and ZBTB18 in the E14 and E16 mouse cortex. (Right) Quantification of ZBTB18-positive neurons in NR4A2-positive cells at E15 and E16. (**E**) Comparison of the NR4A2 expression level in the *Zbtb18* knockout (KO) cortex with that in the wild-type (WT) cortex by microarray analysis. (**F**) Zbtb18-expressed E10-born neurons suppressed NR4A2 expression in the subplate layer. *CAG::GFP*, *CAG::Zbtb18, ^iOn^CAG*∞*IRFP*, and *CAG::hyPBase* are co-electroporated at E10 in the mouse dorsal cortex and fixed at E16. Green signal indicates GFP/Zbtb18, red indicates Nr4a2, and blue indicates DAPI. (**G**) Zbtb18 expression (green) in the cells during cortical development. Scale bars: (a, b) 500 μm. Nuclei are shown in blue using DAPI in all images.

We then examined ZBTB18 expression in the P1 mouse cortex and found it was absent from the *Ctgf*-positive subplate layer (Fig. 4B). scRNA-seq further revealed that *Zbtb18* expression is minimal in SpNs compared with that in other cortical populations (Fig. 4C). The dynamics of this transition were clarified from the developmental time course, showing that ZBTB18 repression initiates at E16, whereas certain SpN markers, such as NR4A2, still co-express with ZBTB18 at E15 (Fig. 4D). Together, these observations suggest that Zbtb18 functions as a key regulator of mSpN fate, whose dynamic downregulation in SpNs coincides with the onset of Nr4a2 expression.

In contrast, Zbtb18 expression is maintained in other early-born neuronal populations, such as those giving rise to layer 4, where Nr4a2 is not induced. Thus, Zbtb18 downregulation acts as a permissive gate that enables Nr4a2 expression and biases early-born neurons toward an mSpN fate. Because TCAs first robustly invade the subplate around E16 (*3*), we determined whether TCA input contributes to Zbtb18 silencing. *Zbtb18* downregulation was attenuated (∼75% Zbtb18-negative) in the subplate cells of *Gbx2* knockout mice lacking TCAs, compared with that in wild-type mice (> 90% Zbtb18-negative), indicating that TCA-dependent input partially drives Zbtb18 repression (Fig. S14).

We then asked whether Zbtb18 directly constrains the mSpN program through Nr4a2. Motif scanning identified consensus *Zbtb18* sites within regulatory regions of *Nr4a2* (Fig. S15). Consistent with this, microarray profiling in the Zbtb18 knockout cortex revealed increased Nr4a2 expression, supporting a de-repression model (Fig. 4E). Conversely, forced Zbtb18 expression abolished NR4A2 in a cell-autonomous manner (Fig. 4F). Across amniotes, Zbtb18 expression diverges: it is selectively repressed in mammalian SpNs but broadly expressed in the pallial neurons of birds and reptiles (Fig. S16). Overall, these findings are consistent with a model in which TCA-dependent input contributes to Zbtb18 repression, thereby de-repressing Nr4a2 and specifying early-born neurons as mSpNs, reflecting a repurposing of an ancestral input-neuron program (Fig. 4G).

## Discussion

Previous studies have suggested heterogeneity and multiple developmental origins of SpNs (*13, 24*). Building on these observations, our findings provide direct evidence for a dual-origin framework in which mammalian SpNs comprise two evolutionarily distinct lineages: an ancestral, medial pallium-derived subtype (aSpNs) and a mammalian-specific, dorsal pallium-derived subtype (mSpNs) arising from early-born cortical neurons. This framework provides a unifying explanation for why canonical SpN-associated markers (e.g., Ctgf, Nr4a2, Moxd1) are observed outside mammals and why marker presence alone cannot establish homology in sauropsid pallia. We further identify a mechanism underlying this divergence, in which temporal repression of Zbtb18 gates mSpN specification during development (Fig. 5). The identification of a medial pallium-derived SpN population parallels the multi-source origin described for Cajal–Retzius cells (*25*), suggesting that multi-origin developmental logic is a general feature of early cortical neurogenesis.

**Fig. 5.**
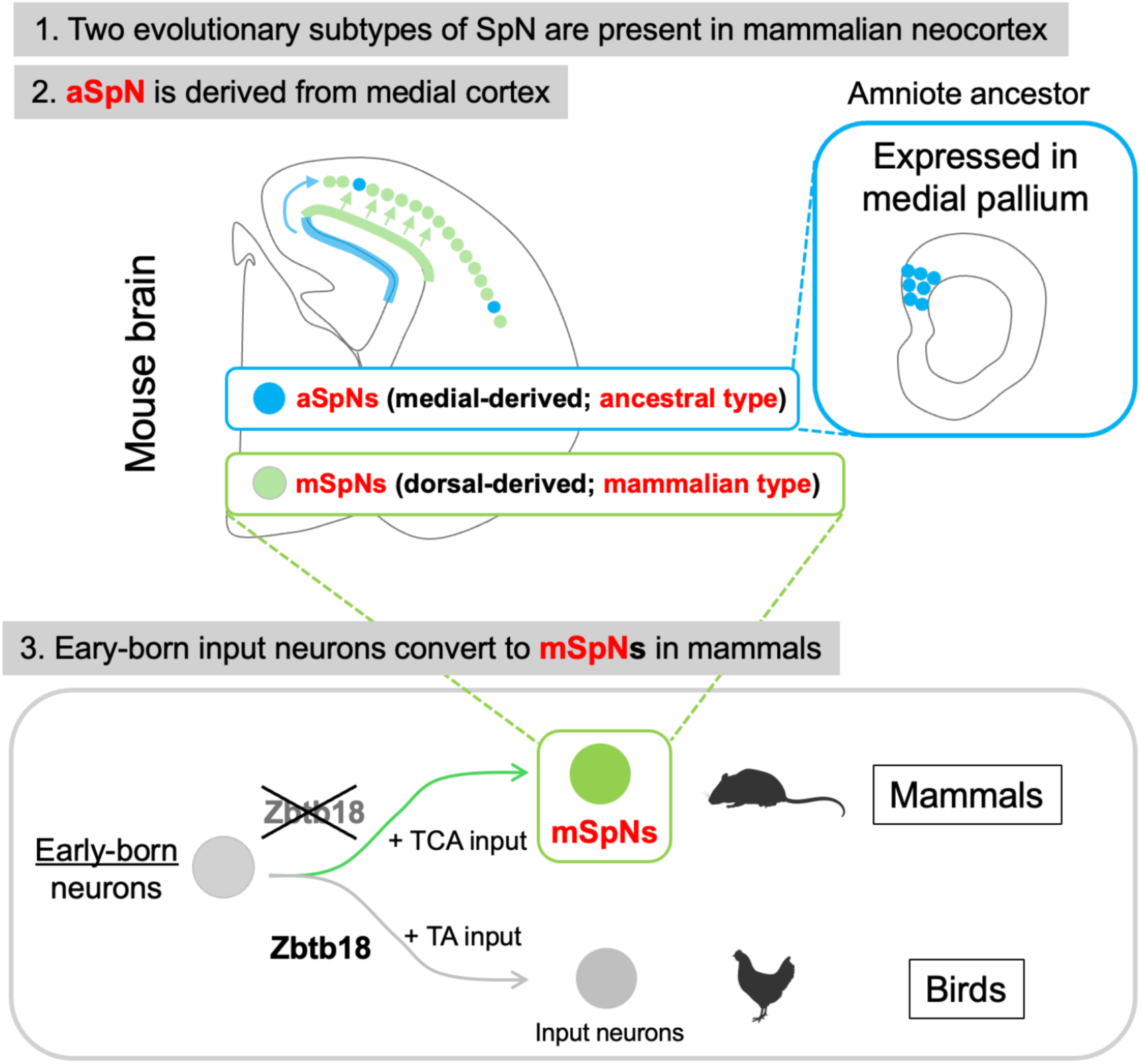
Summary of the key findings in this study.

Comparative analyses suggest that early-born pallial neurons across amniotes adopt an input-neuron-like program. In non-mammalian species, such as chicks and reptiles, these neurons differentiate into thalamorecipient input neurons. In mammals, this ancestral program is repurposed: early-born neurons adopt a transient subplate identity, whereas later-born layer 4 neurons establish the canonical thalamic input layer.

This evolutionary redistribution of input functions provides a framework for the emergence of the six-layered neocortex. By inserting a transient, activity-sensitive layer ahead of layer 4, mammals may have gained both temporal flexibility and developmental robustness during thalamocortical circuit assembly. This framework also provides a potential link between subplate evolution and neocortical expansion. As cortical size and connectivity increased during mammalian evolution, the need to coordinate long-range thalamocortical interactions during early development would have imposed substantial constraints (*3, 5*). The transient deployment of an input-like subplate layer may have provided a developmental solution to these constraints, enabling early network coordination (*26*) while preserving the integrity of the emerging laminar architecture. Such a mechanism may have been particularly important in supporting the emergence of large-scale, highly interconnected cortical networks characteristic of mammalian brains.

Our data identify Zbtb18 as a central regulator of this repurposing. Zbtb18 is broadly expressed in the amniote pallium but selectively repressed in SpNs, and its downregulation coincides with TCA invasion. This downregulation is attenuated in the TCA-deficient cortex and is associated with the de-repression of Nr4a2, supporting a model in which activity-dependent signals enable the emergence of a subplate identity. In this framework, Zbtb18 functions as a permissive “fate gate” that regulates access to alternative developmental trajectories in early-born neurons. This suggests that new cortical cell types arise not only through the emergence of new gene programs but also through changes in how existing developmental programs are deployed.

Recent work has identified ZBTB18 as a key regulator of projection neuron diversification in the mammalian neocortex through mammalian-specific cis-regulatory elements (*27*). Our findings extend this framework by revealing a complementary mechanism: in addition to activating projection neuron programs, ZBTB18 contributes to cortical diversification through selective repression. Specifically, the downregulation of Zbtb18 releases a latent developmental trajectory, enabling early-born neurons to adopt a subplate identity via Nr4a2 de-repression. Hence, Zbtb18 may function not only as a developmental regulator but also as a candidate evolutionary switch that enables the repurposing of an ancestral input-neuron program into a mammalian-specific subplate identity.

Nonetheless, the upstream signals that regulate Zbtb18 downregulation during subplate development have yet to be identified. The subplate is characterized by a specialized extracellular matrix (ECM) niche enriched in components such as neurocan, versican, and syndecans, which regulate neuronal polarity, migration, survival, and differentiation. TGF-β, present in an inactive form within the subplate ECM, is activated by the protease Adamts2 secreted from migrating neurons, thereby promoting a switch in neuronal migration modes within the subplate (*28*). This ECM also serves as a reservoir for multiple signaling molecules and contributes to diverse signaling events in and around the subplate. Although the mechanism underlying Zbtb18 downregulation in SpNs remains unclear, TGF-β signaling represents a plausible upstream candidate and may constitute a conserved pathway regulating Zbtb18 dynamics across amniotes.

Ultimately, our cross-species integration is subject to limitations, including embryonic sampling in sauropsids, potential transcript capture biases, and constraints in ortholog mapping. Future studies combining lineage tracing with temporally resolved multi-omics and targeted perturbations of Zbtb18, thalamocortical input, and local niche signals are needed to establish causality at each step.

Despite these limitations, our findings support a dual-origin model in which the mammalian subplate emerges through activity-dependent rewiring of an ancestral input-neuron program via Zbtb18 silencing. By transiently deploying an input-like layer during early development, this mechanism may have provided a solution to the challenges associated with coordinating long-range thalamocortical interactions as the cortical size and connectivity increased. More broadly, our results suggest that the evolution of mammalian cortical circuits relies not only on the emergence of new cell types but also on the temporal redeployment of conserved developmental programs, revealing a general principle by which complex brain architectures can develop, consistent with the idea that new cell types can arise through evolutionary modification and the diversification of regulatory programs (*29*).

## Acknowledgments

We thank all members of the developmental neuroscience project at TMiMs for their helpful comments on this project.

## Author contributions

T.K. and C.O.-M. designed the research. T.K. performed the main experiments and analyzed the data. Y.H. performed scRNA-seq analyses, including preprocessing, clustering, and cross-species integration. Y.H., I.A., and H.A. analyzed the scRNA-seq and Visium datasets. R.K. performed chick experiments. Y.N. performed *in utero* electroporation into the thalamus. S.G.-S. and R.W. performed CAGE experiments. H.H. generated *Gbx2* knockout mice. K.N., H.K., and C.O.-M. provided supervision and reviewed the manuscript. T.K. and C.O.-M. wrote the paper. All authors reviewed and approved the final manuscript.

## Data and materials availability

All data are available in the main text or supporting information

## Materials and Methods

### Animals

All animals were treated in accordance with the Tokyo Metropolitan Institute of Medical Science Animals Care and Use Committee guidelines. Pregnant ICR mice were purchased from Japan SLC Inc. (Hamamatsu, Japan) and used for *in utero* electroporation, birthdate analysis, and immunofluorescence. *Gbx2* knockout (KO) mice were kindly provided by Drs. Kazunori Nakajima, Satoshi Yoshinaga, and Hiroyuki Hasegawa (Keio University). *Gbx2* KO mice were produced using the i-gonad method (*1*). Fertilized chick eggs were purchased from Yamagishi (Mie, Japan). Chick eggs were incubated in a humidified incubator at 37°C until collection. Fertilized turtle eggs were purchased from Yamato Yoshoku (Saga, Japan). Turtle eggs were incubated in a humidified incubator at 30°C until collection. Fertilized Madagascar ground gecko (*Paroedura picta*) eggs were kindly provided by Drs. Hiroshi Kiyonari and Takeya Abe (RIKEN, BDR).

### Antibodies

The primary antibodies used for immunostaining were rabbit anti-FOG2 (ZFPM2) (1:250; Santa Cruz), rat anti-Ctip2 (1:1000, Abcam), rabbit anti-Tbr1 (1:1000, Abcam), rabbit anti-Tle4 (1:250; Santa Cruz), chick anti-GFP (1:1000, Abcam), goat anti-Ctgf (1:500, Santa Cruz), rabbit anti-BF1 (1:1000, TaKaRa), goat anti-Nr4a2 (1:500, R&D), rabbit anti-Zbtb18 (1:1000, gift from Dr. H. Okado), and rabbit anti-MAP2 (1:500, CST). The secondary antibodies were Alexa Fluor 488-conjugated donkey anti-chicken IgY (IgG) (703-545-155, Jackson ImmunoResearch), Cy5-conjugated donkey anti-rabbit IgG (711-175-152, Jackson ImmunoResearch), Alexa Fluor 546-conjugated donkey anti-rabbit IgG (A10040, ThermoFisher Scientific), and Cy5-conjugated donkey anti-mouse IgG (715-175-150, Jackson ImmunoResearch). The antibodies were used at a 1:500 dilution unless otherwise noted.

### Immunostaining

For the mouse cortex, embryos fixed for 1 h in 4% paraformaldehyde (PFA) were equilibrated in 30% sucrose and embedded in TissueTek (Sakura), frozen on dry ice, and stored at 80°C prior to cryostat sectioning (Microm HM560, 20-μm sections). After equilibration at room temperature, sections were washed in PBS before blocking in PBS-0.1% Triton-10% normal donkey serum (NDS) and incubating overnight with primary antibodies in PBS-0.1% Triton-1% NDS. Following PBS washes, slides were incubated for 1 h with secondary antibodies in the above buffer, washed, and mounted with Mowiol/DABCO.

### Neuronal Birth Dating

For the birthdate analysis, the thymidine analog EdU was injected to label newborn cells.

Eggs: A small hole was made in the top of the eggshell using dissection scissors, the embryo was exposed by removing the surrounding membranes, and 5 mL of albumen was removed with a syringe to make sufficient space for the injected volume. EdU (5 μg/μL) was injected into the cerebrum by mouth pipetting into the ventricle. After applying 5 μg/mL gentamicin sulfate/PBS heated to 37°C into the open eggshell, the hole in the eggshell was sealed with tape. Eggs were returned and incubated until E14. Embryos were fixed with 4% PFA in PBS.

Mice: Pregnant dams were given intraperitoneal injections with single pulses of EdU at E11 or E12 (50 mg/kg body weight). Embryos were harvested at P0, perfused, and fixed in 4% PFA for 1 hr. For EdU-double labeling with RNAscope, *Ctgf* mRNA was first detected using the RNAscope Multiplex Fluorescent V2 kit, followed by EdU visualization using the Click-iT EdU kit (Invitrogen). Similarly, for EdU-double immunohistochemistry, EdU visualization using the Click-iT EdU kit followed the detection of the ZFPM2, CTIP2, TBR1, or TLE4 antigens.

### RNAscope

Coronal sections of wild-type brains (mice, chicks, turtles, and geckos) were used for this study. For *Ctgf* and *Eag2* mRNA detection, we used the RNAscope Multiplex Fluorescent V2 kit (Advanced Cell Diagnostics) and fluorescein, Cy3, and Cy5 TSA fluorophore (Akoya Biosciences) according to the manufacturer’s protocol with minor modifications. Briefly, cortical sections (20 μm thick) were obtained using a cryostat, and the slides were subsequently stored at –80°C until use. Coronal brain sections were baked at 60°C for 30 min, fixed with 4% paraformaldehyde in PBS on ice, washed with 100% ethanol, and then treated with hydrogen peroxide for 10 min at room temperature. After washing with water, the sections were boiled for 5 min in a target retrieval buffer (Advanced Cell Diagnostics). Following target retrieval, sections were washed sequentially with water and 100% ethanol, and then left to dry overnight. Next, the sections were treated with the protease for 10 min at 40°C and exposed to the *CTGF* and *EAG2* probes (Advanced Cell Diagnostics) for 2 h at 40°C before signal amplification. The fluorescence signal was further amplified using the TSA plus system (Akoya Biosciences) before detection. Sections were finally washed and treated with DAPI for 2 min before mounting in FluoroKEEPER Antifade Reagent (Nakalai Tesque).

### Mouse and chick embryonic electroporation

*In utero* and *in ovo* electroporation were performed in mouse and chick embryos as previously described (*2*, *3*).

#### Mice

*In utero* electroporation was performed at embryonic day (E)10.5 or E11.5. A DNA mixture containing 0.5–1 μg/μL of iOn vector, 0.5–1.2 μg/μL of a non-integrative control plasmid, and 0.2 μg/μL of *CAG::hyPBase* plasmid supplemented with Fast Green dye was injected into one lateral ventricle using a glass capillary pipette. The heads of the embryos in the uterus were placed between a tweezer-type electrode 1-mm in diameter disk electrode (LF650P1, BEX), and then four electric pulses (50V, 50 ms in duration at intervals of 950 ms) were delivered using a CUY21E electroporator (BEX). Embryos were allowed to develop until the indicated stages. Tissues were fixed in 4% PFA for 1 h and then transferred to PBS containing 0.1% NaN₃.

#### Chicks

*In ovo* electroporation was performed at E4. After removal of the albumen, a window was opened at the top of the eggshell. DNA solution was injected into the lateral ventricle using a mouth pipette. Electrodes were positioned on the head’s surface, and square pulses (15 V, 50 ms duration with 950 ms intervals) were applied using a pulse generator (GEB15, BEX). The eggshell was sealed with tape, and embryos were incubated until E10 or E14. Embryos were then fixed in 4% PFA in PBS.

### CAGE-seq

The quality of total RNA was assessed using an Agilent 2100 Bioanalyzer (Agilent Technologies, Santa Clara, CA, USA) to ensure that the RNA integrity number was over 7.0. CAGE libraries were prepared using a CAGE Library preparation kit (DNAform, Kanagawa, Japan). Briefly, cDNA was synthesized from total RNA using random primers. The ribose diols in the 5′ cap structures of RNA were oxidized and then biotinylated. The biotinylated RNA/cDNAs were selected by employing streptavidin beads (a process known as cap-trapping). RNA was digested with RNaseONE/H, adaptors were ligated to both ends of cDNA, and CAGE libraries of double-stranded cDNA were constructed. The CAGE libraries were validated for quality and size distribution using an Agilent 2100 Bioanalyzer and sequenced to an average depth of approximately 50 million reads using a NextSeq500 system (Illumina, San Diego, CA, USA).

### Brain dissociation and single-cell preparation

Dorsal pallial tissue was dissected from the brains of chicks and turtles and immediately transferred to homogenization tubes containing 1 mL of TRIzol reagent (ThermoFisher Scientific). Following homogenization, total RNA was extracted by phenol-chloroform phase separation. RNA was precipitated with isopropanol, washed, and resuspended in RNase-free water. Samples were stored at −80°C until further analysis. RNA concentration and purity were assessed using a NanoDrop spectrophotometer. The extracted RNA was used for CAGE analysis as described above, and for RNA-seq performed in Novogene using a poly-A selection protocol with 150 bp paired-end reads sequencing.

For the developmental scRNA-seq dataset, pallial cells were isolated from E14 chick and P2 turtle brains. Tissue pieces were dissociated with papain for 25 min at 32°C. After four washes with culture medium, the dissociated cells were resuspended in 200 μL of culture medium, passed through a membrane filter, and collected in a 5-mL FACS tube. Propidium iodide (10–20 μL of a 10 μg/mL solution) was added to the cell suspension, and propidium iodide-negative viable cells were isolated by FACS using a SONY SH800 cell sorter. After sorting and concentration, cells were resuspended in PBS containing 2% FBS at approximately 2,000 cells/μL. The cell suspension was loaded onto a 10x Chromium Chip G with a targeted recovery of 8,000 cells for GEM generation and cell barcoding. Single-cell RNA-seq libraries were prepared using the 10x Chromium Next GEM Single Cell 5′ Reagent kit v2 (Dual Index).

### Spatial gene expression profiling

Brains were collected from mouse embryos at E17 and P0, and from chick embryos at E14, and embedded in O.C.T. compound. The embedded brains were stored at −80°C until sectioning. Frozen sections (10 μm thick) were prepared according to the manufacturer’s protocol. Sections were mounted onto Visium Spatial Gene Expression Slides (10x Genomics, USA) and processed using the Visium Spatial Gene Expression Reagent kit (10x Genomics, USA) following the manufacturer’s instructions. Hematoxylin and eosin (H&E) staining and fluorescence images were acquired as tiled mosaics using a Keyence BZ-X810 microscope (Keyence, Japan). Libraries were prepared according to the Visium protocol and sequenced by on a NovaSeq (S1) platform (Illumina, USA) with 150-bp paired-end reads.

### Data processing of spatial gene expression

FASTQ files from the chick brain Visium sample were processed using Space Ranger v1.3.1 (10x Genomics, USA) with default parameters and the NCBI RefSeq gene annotations for the galGal6 genome assembly, generating spatial gene expression matrices for individual spots. The gene expression dataset was analyzed using Seurat v4.4.0 (4). Read count matrices were normalized and integrated using the anchor-based integration workflow implemented in the Seurat FindIntegrationAnchors and IntegrateData functions (Stuart et al., 2019).

Spots from the three samples were clustered based on gene expression profiles using the shared nearest neighbor (SNN) approach with the Seurat FindNeighbors and FindClusters functions at a resolution of 0.5, resulting in eight clusters. Uniform Manifold Approximation and Projection (UMAP) was performed for dimensionality reduction using the Seurat RunUMAP function. Two-dimensional UMAP plots were visualized using the Seurat DimPlot function and the Loupe Browser application. Marker genes for each cluster were identified using the Seurat FindMarkers function.

Processed Visium datasets from E17 and P0 mouse brains were obtained from Hara et al. (2025) (5). Spatial gene expression patterns were visualized using Loupe Browser v8.0.0 (10x Genomics, USA).

### Data processing of gene expression of scRNA-seq

For scRNA-seq data processing, the mouse genome assembly mm10 and gene annotation (refdata-gex-mm10-2020-A) were obtained from the 10x Genomics website. The Chinese softshell turtle and chicken genome assemblies, PelSin_1.0 and galGal6, as well as their gene annotations, were retrieved from Ensembl (release 109 for turtle and release 104 for chicken). The turtle gene annotation was further refined by incorporating bulk RNA-seq and CAGE-seq data to accurately count reads mapped to 5’ UTR ends. These bulk RNA-seq and CAGE-seq reads were obtained from samples of the same tissue and developmental stage as those used for scRNA-seq. The sequence reads were mapped to the genome assembly PelSin_1.0 using STAR v2.7.10b (*6*) in a splicing-aware manner. A transcriptome-oriented gene annotation was created using these mapping data by employing StringTie v2.1.4 (*7*), guided by the Ensembl release 109 gene annotation, and the improved annotation was incorporated into the existing one. Finally, the assemblies and annotations were prepared for the Cell Ranger pipeline using the “cellranger mkref” command implemented in 10x Genomics Cell Ranger v7.1.0 (*8*).

Five-prime scRNA-seq reads of the chicken and turtle were generated as described in the previous section, and 3’ scRNA-seq reads of the E18.5 mice cortex were retrieved from NCBI GEO (GSM4635078 and GSM4635079) (*9*). These scRNA-seq reads were processed using the “cellranger count” command in 10x Genomics Cell Ranger v7.1.0 with default parameters, which included auto-detection of the types of sequencing libraries, referring to the aforementioned genome assemblies and gene annotations of individual species. This process generated the count matrices of the genes for the individual cells. All subsequent scRNA-seq analyses were performed using Seurat v4.9.9.9045 with default parameters, except as noted for the parameters of certain functions (*10*). The count matrices were normalized by employing the SCTransform approach (*11*) after discarding the cells with fewer than 700 UMI counts or more than 15% mitochondrial contamination. The gene expression profiles of the individual cells were visually inspected using 10x Genomics Loupe Browser v8.0.0 by converting the Seurat object into a Loupe file using the loupeR package v1.0.0 in R (https://github.com/10XGenomics/loupeR).

### Cross-species integration of scRNA-seq data

To accommodate the count matrices of scRNA-seq data across chicks, turtles, and mice, we created subsets of the matrices that contained one-to-one orthologs retained by the three species. The one-to-one orthologous relationships among the three species were retrieved from Ensembl release 98, followed by the replacement of Ensembl Gene ID with those in the newer releases used for read counting. The gene symbols of the chicken and turtle were replaced with those of the mouse, allowing us to consistently integrate gene count matrices across multiple species. Cells with low UMI counts or high mitochondrial contamination ratios were discarded from the matrices in the same way described above, and subsequently, the matrices consisting of one-to-one orthologs were created. The Seurat objects of the three species containing the one-to-one ortholog subsets were integrated by employing the anchor-based approach (*12*), followed by NNI-based clustering of the cells with the resolution parameter set at 0.25. The resulting 16 clusters were manually annotated, referring to the marker genes listed in the marker gene subsection.

Glutamatergic neurons were identified from the cross-species integration data by selecting cells from clusters 1, 2, 3, 9, and 10 (Fig. S3) and further discarding potential GABAergic neurons expressing (positive count values) either Gad1 or Gad2 genes. These glutamatergic neurons from the three species were again subjected to cross-species integration by implementing the anchor-based approach and NNI-based clustering with the resolution parameter set at 0.35. In line with the clusters with fewer cell numbers, the first to sixteenth dimensions for the canonical correlation analysis were employed with k.filter=20 and the k.score=15 for the Seurat FindIntegrationAnchors function and with k.weight=15 for the Seurat IntegrateData function. Cells in the cluster 8 (gCl8) of the glutamatergic neurons, the candidates for subplate neurons and their homologous cells, were reanalyzed with an integration by employing the reciprocal PCA (RPCA) approach. This integration was first performed using the chicken and mouse cells, because gCl8 contained only three turtle cells. The integration employed the first to twentieth dimensions with k.filter=80 for the Seurat FindIntegrationAnchors function and with k.weight=20 for the Seurat IntegrateData function. The turtle cells were then projected onto the reference UMAP space of chicken and mouse cells. The projected PCA embeddings of the turtle cells were transformed into this UMAP space using umap_transform (n_epochs = 55) implemented in the uwot v0.1.16 package (https://cran.r-project.org/web/packages/uwot/). The resulting UMAP coordinates were combined using Seurat.

### Integration with single-cell RNA-seq

The scRNA-seq expression values of a mouse developing brain were retrieved from mousebrain.org (http://mousebrain.org/development/downloads.html). The dataset, which was stored in the loom format, was converted into the ReadVelocity function implemented in the SeuratWrapper R package and the as.Seurat function. We extracted the expression profiles of forebrain neural cells of stage E17 from the dataset to integrate them into the spatial expression profiles of the E17 forebrain slice. The cluster annotation of single cells was used for the following analysis.

The expression values of the spots for the E17 slice were normalized by employing the SCTransform approach (Hafemeister and Satija 2019) with the Seurat SCTransform function. These spots were then clustered based on gene expression profiles with the Seurat FindNeighbors and FindClusters functions, and dimension reduction of gene expression profiles was performed with the Seurat RunPCA and RunUMAP functions. The expression values of E17 forebrain single cells were processed similarly. Integration of the scRNA-seq dataset into the spatial transcriptome one was performed with the Seurat FindTransferAnchors function (*10*), employing an anchor-based approach by querying the spatial gene expression profiles and referring to single-cell gene expression profiles. The integrated dataset was further processed with the Seurat TransferData function, which yielded a prediction score—a confidential index of cell type prediction—of the individual single-cell clusters for every spot. We chose the single-cell clusters that exhibited high prediction scores in the spots of Visium clusters 21 (choroid plexus) and 22 (claustrum). The prediction scores were visualized by overlaying them onto the Visium spot with the Seurat SpatialFeaturePlot function (Figs. 4d and S2a). The selected single-cell clusters were visualized as a UMAP plot following the aforementioned procedures, and the clusters were grouped by visual inspection based on the UMAP plot. Among these groups, we searched for marker genes more highly expressed in one group than the others using the Seurat FindMarkers function.

### Marker genes

Cell types were annotated using canonical marker genes: *Gad2* for GABAergic neurons, *Neurod2* for glutamatergic neurons, *Nr4a2* and *Ctgf* for subplate-associated populations, *Olig2* and *Pdgfra* for oligodendrocyte precursor cells (OPCs), *Plp1* for mature oligodendrocytes, *Sox9* for astrocyte progenitors, and *Slc1a3* and *Gfap* for mature astrocytes.

### Data availability

Sequence reads were deposited in the NCBI Short Read Archive (*4*) under BioProject accession number PRJNA1465250, and the Visium data with SpaceRanger and slide images were deposited in the DDBJ under the DataSets accession number SSUB020519 and SSUB024228.

---

**Fig. S1.**
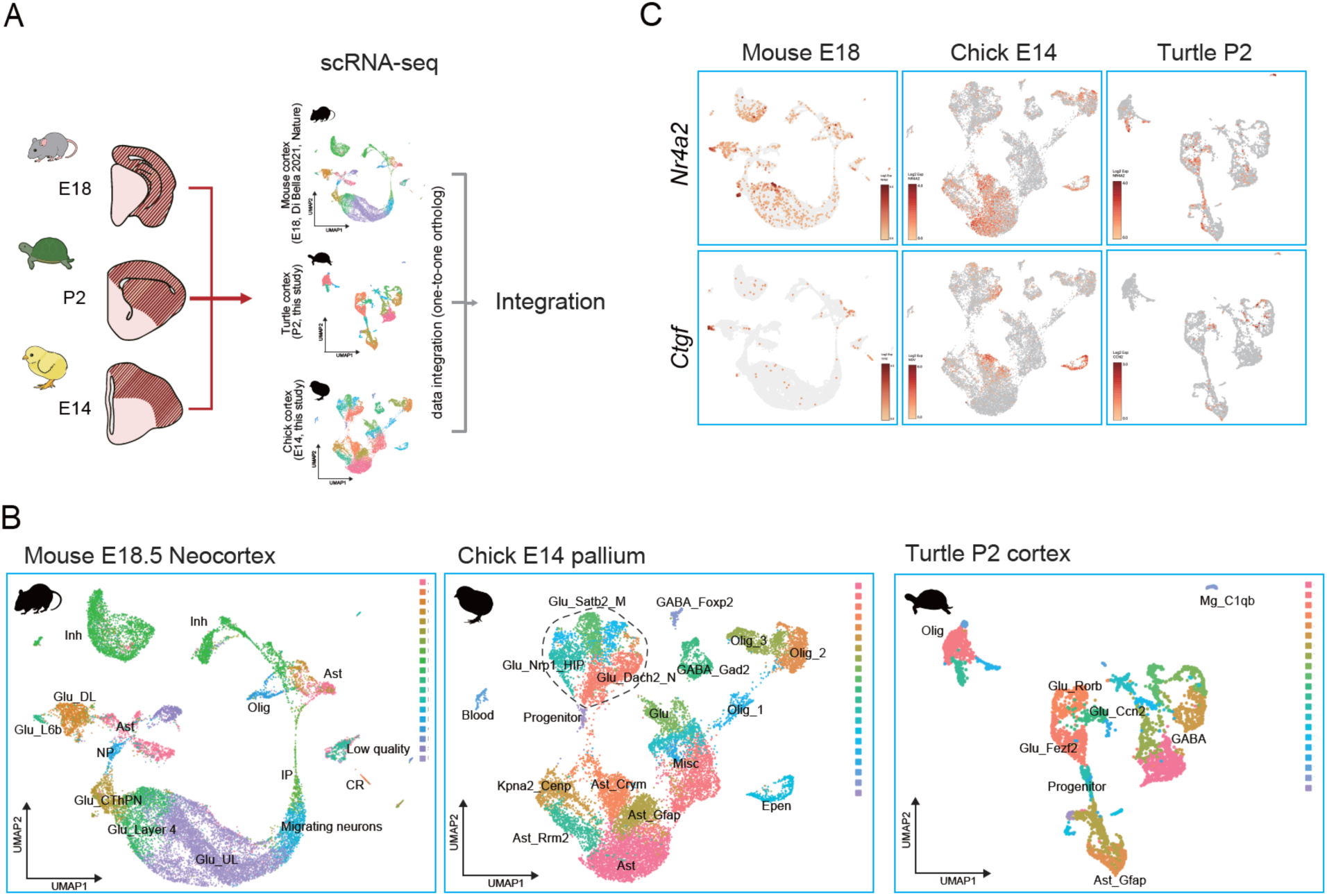
Cross-species integration of amniote scRNA-seq datasets. (**A**) Workflow for scRNA-seq integration across three amniote brains: mouse E18, turtle P2, and chick E14. (**B**) UMAP projections shown separately for each species. Clustering was performed within each species. (**C**) Expression of SpN markers (*Nr4a2* and *Ctgf*) on species-specific UMAPs.

**Fig. S2.**
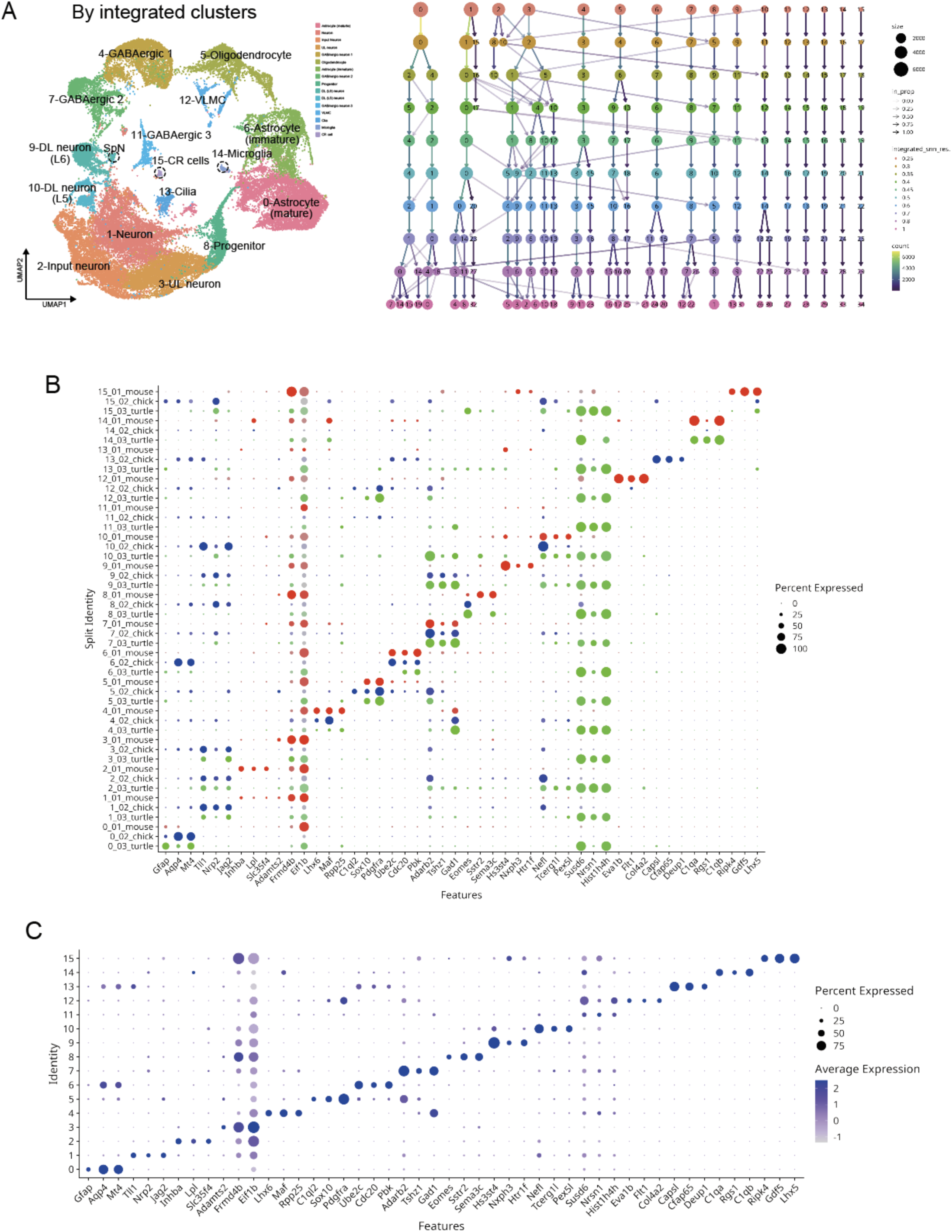
Cross-species integration across amniote scRNA-seq datasets. (**A**) Left: UMAP of the integrated dataset colored by integrated clusters and annotated with major cell classes. Right: Cluster tree across Seurat clustering resolutions (integrated_snn_res.) showing relationships and stability of integrated clusters; node size reflects cell number and edges indicate transitions between resolutions. (**B**) Dot plot of selected marker genes across integrated clusters split by species (mouse, chick, and turtle). Dot size indicates the percentage of cells expressing each gene, and color indicates scaled average expression. (**C**) Dot plot of the same marker genes across integrated clusters in the combined (species-integrated) dataset. Dot size indicates the percentage of cells expressing each gene, and color indicates scaled average expression.

**Fig. S3.**
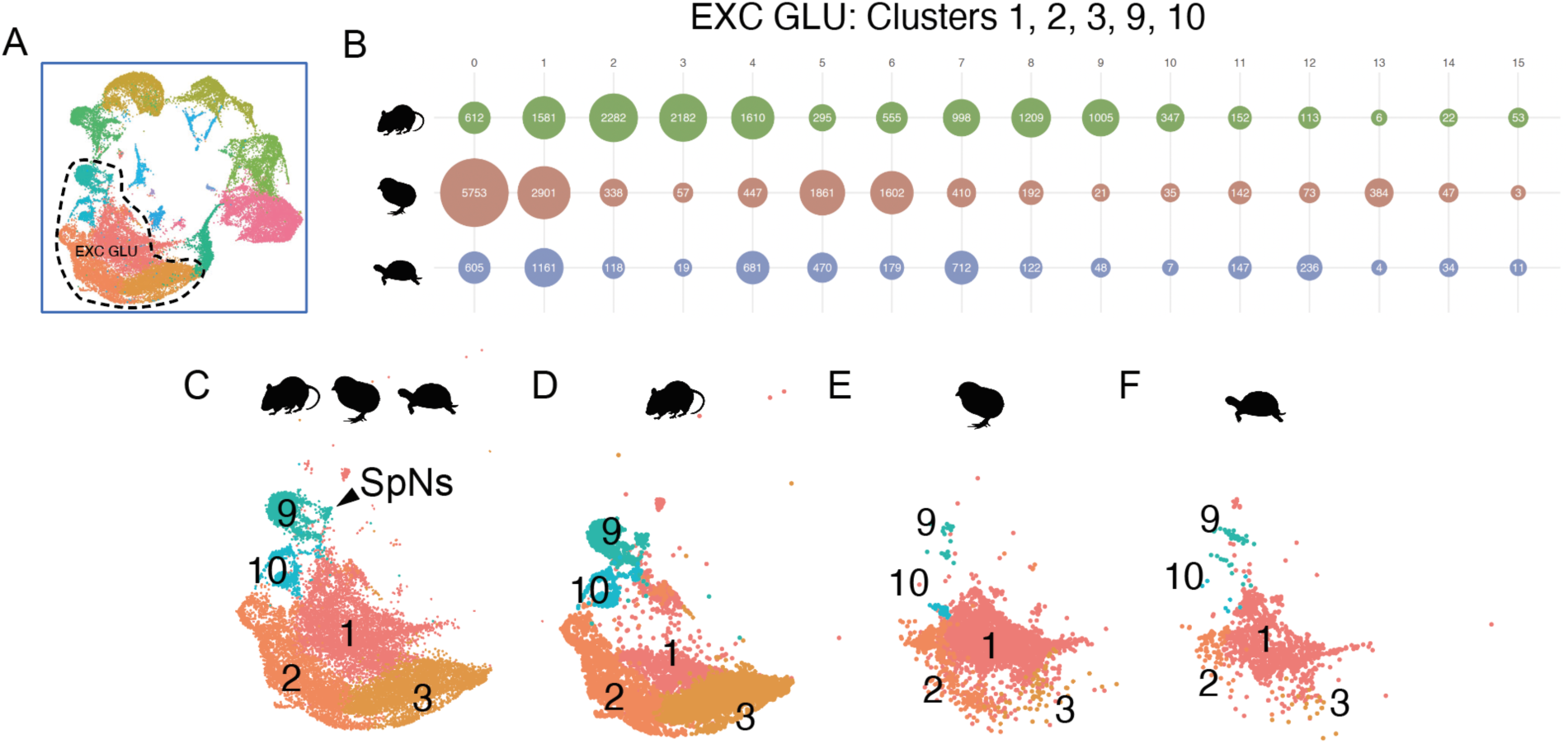
Species composition and UMAP distribution of excitatory glutamatergic (EXC_GLU) clusters in the integrated amniote scRNA-seq atlas. (**A**) Overview of the integrated UMAP highlighting the EXC_GLU compartment (dashed outline). (**B**) Cell numbers per integrated cluster for each species within the EXC_GLU compartment. Circle size reflects the number of cells, and numbers indicate cell counts. (**C–F**) UMAPs of EXC_GLU cells colored by integrated cluster identity. (C) All species combined (mouse + chick + turtle). (D) Mouse cells only. (E) Chick cells only. (F) Turtle cells only. Integrated EXC_GLU clusters 1, 2, 3, 9, and 10 are shown.

**Fig. S4.**
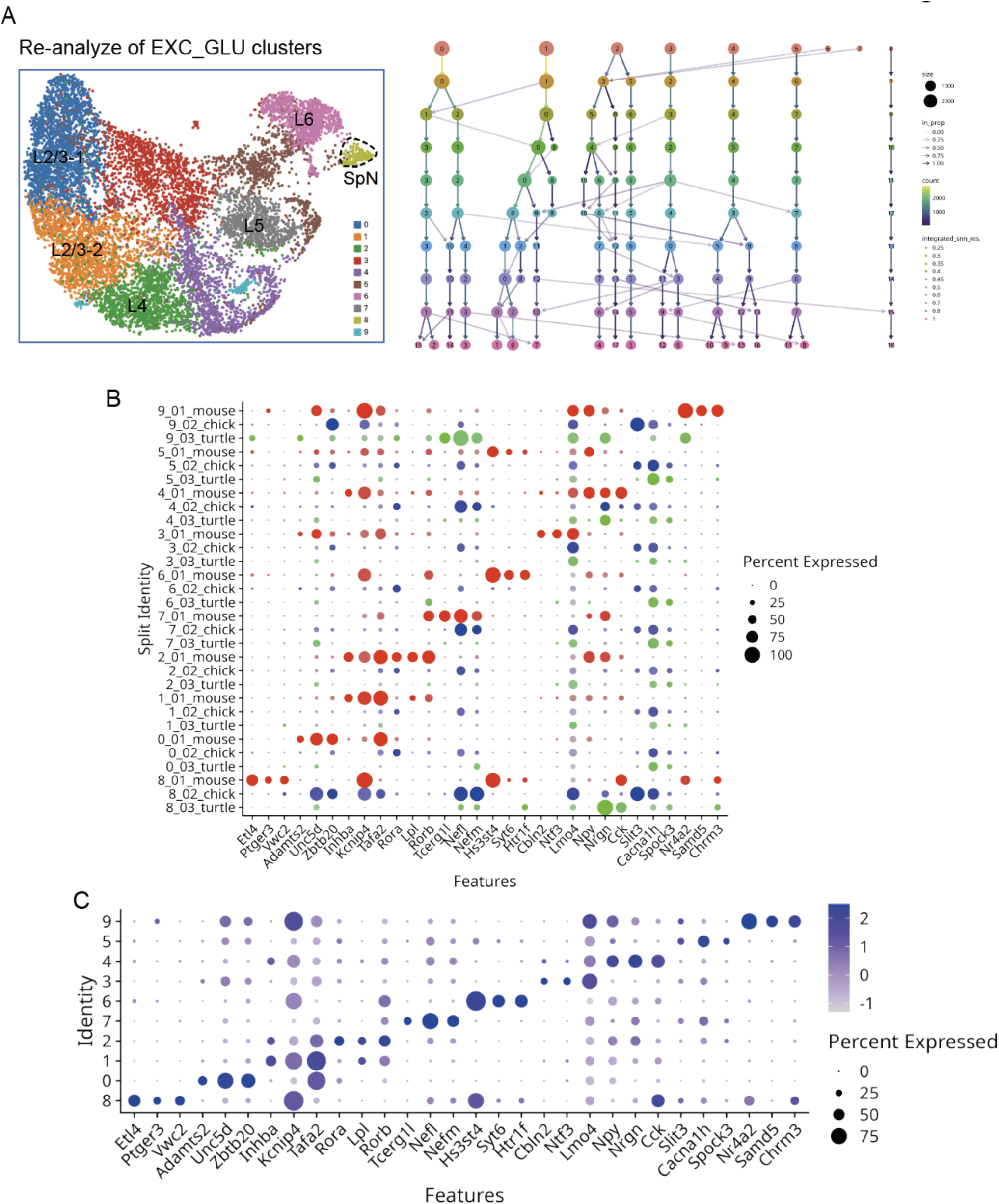
Re-analysis of EXC_GLU populations resolves cortical-layer-like subclusters and their marker gene profiles across species. (**A**) Left: UMAP of re-analyzed excitatory glutamatergic neurons (EXC_GLU) from the integrated amniote scRNA-seq dataset, colored by subclusters and annotated with layer-like domains (L2/3, L4, L5, L6) and an SpN-enriched region. Right: Cluster tree across Seurat clustering resolutions (integrated_snn_res.) illustrating relationships and stability among EXC_GLU subclusters; node size reflects cell number. (**B**) Dot plot of selected marker genes across EXC_GLU subclusters, split by species (mouse, chick, and turtle). Dot size indicates the percentage of cells expressing each gene, and color indicates scaled average expression. (**C**) Dot plot of the same marker genes across EXC_GLU subclusters in the combined integrated dataset. Dot size indicates the percentage of cells expressing each gene, and color indicates scaled average expression.

**Fig. S5.**
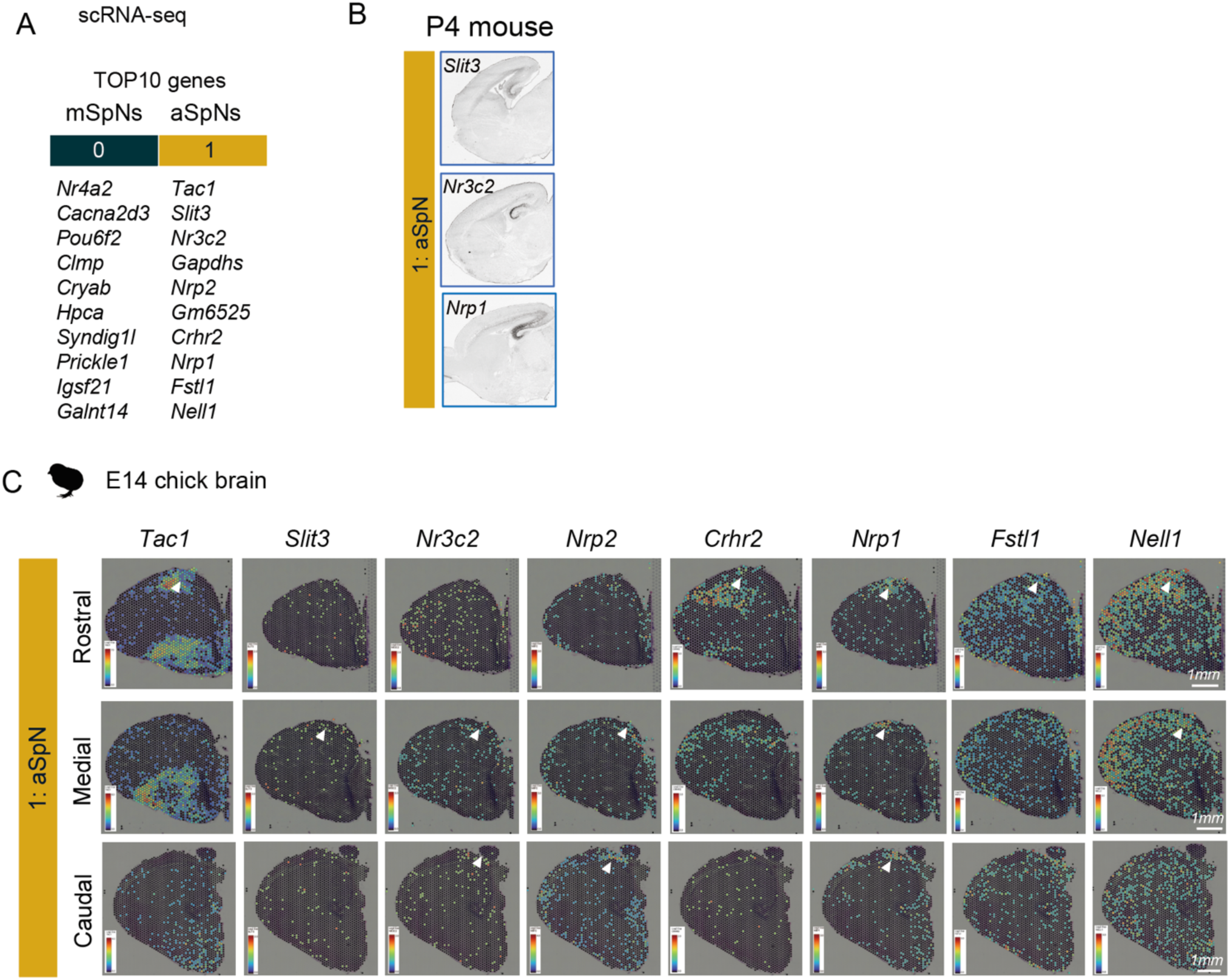
Cross-species candidate markers and spatial expression patterns of EXC_GLU subclusters. (**A**) Top-ranked marker genes for mouse EXC_GLU subclusters corresponding to mSpNs and aSpNs. Cluster 0 and cluster 1 are indicated by color bars. (**B**) *In situ* hybridization images from the P4 mouse brain showing representative aSpN-enriched candidate genes, obtained from the Allen Brain Atlas. (**C**) Spatial feature plots of selected aSpN candidate genes in the E14 chick pallium using Visium. Expression patterns are shown across rostral–caudal levels. Arrowheads indicate regions with enriched expression. Color scales indicate normalized expression levels.

**Fig. S6.**
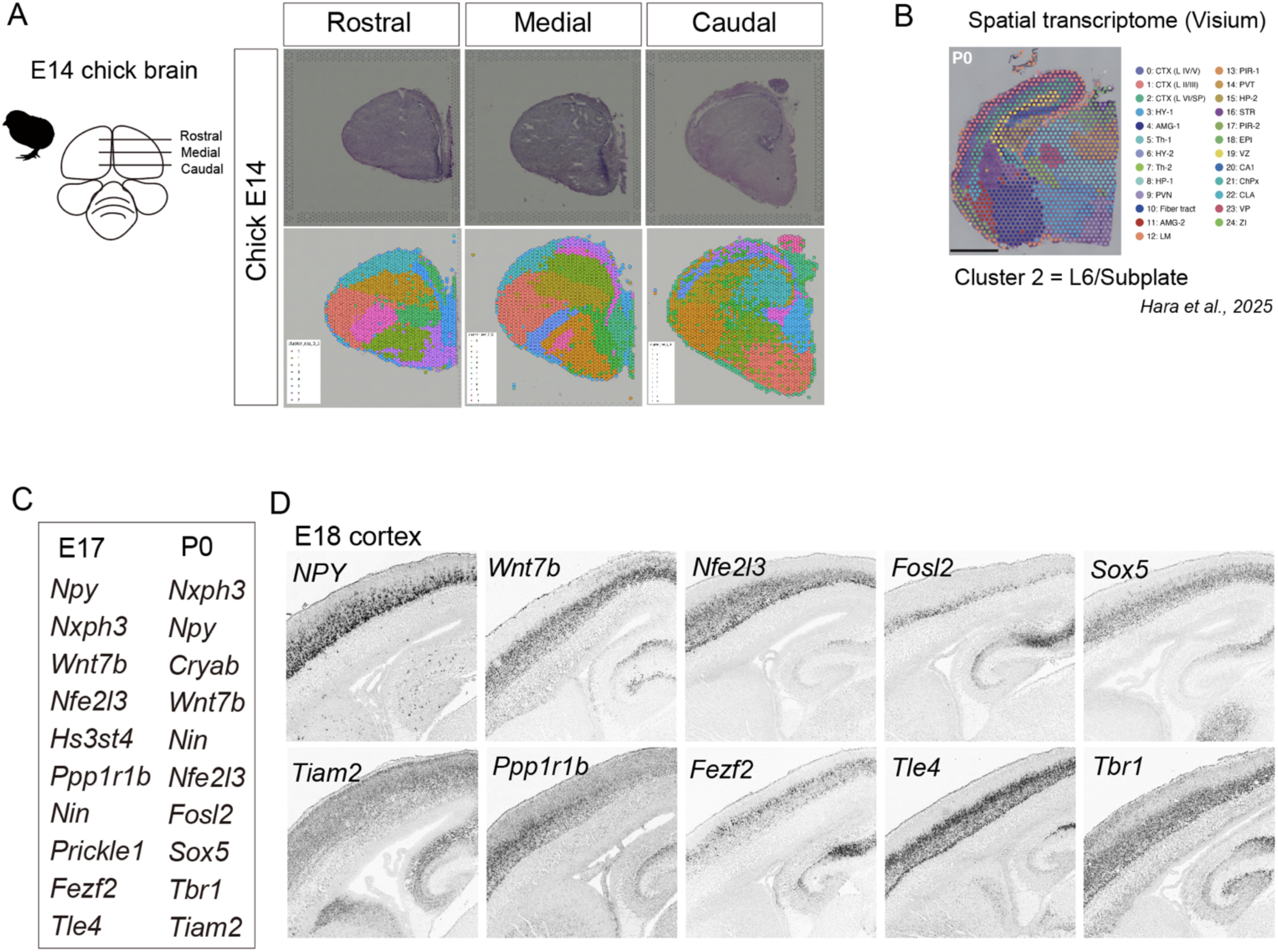
Spatial transcriptomics in mouse and chick embryonic brains. (**A**) Spatial expression mapping of mouse subplate neuron–associated genes onto the chick E14 pallium using Visium. Three rostrocaudal levels of the chick brain are shown: rostral, medial, and caudal sections. Upper panels show histological images, and lower panels show spatially resolved spot clusters overlaid on the corresponding sections across the chick pallium. (**B**) Spatial transcriptomic clustering of the mouse P0 cortex based on Visium data from Hara et al. (2025). Cluster 2 corresponds to the L6/subplate-enriched domain. (**C**) Summary of representative genes enriched in the mouse subplate/L6-related cluster at E17 and P0. (**D**) *In situ* hybridization images showing the expression patterns of representative subplate/L6-associated genes in the mouse E18 cortex, obtained from the Allen Brain Atlas. These genes are enriched in deep cortical layers and/or subplate-related domains in the developing mouse cortex.

**Fig. S7.**
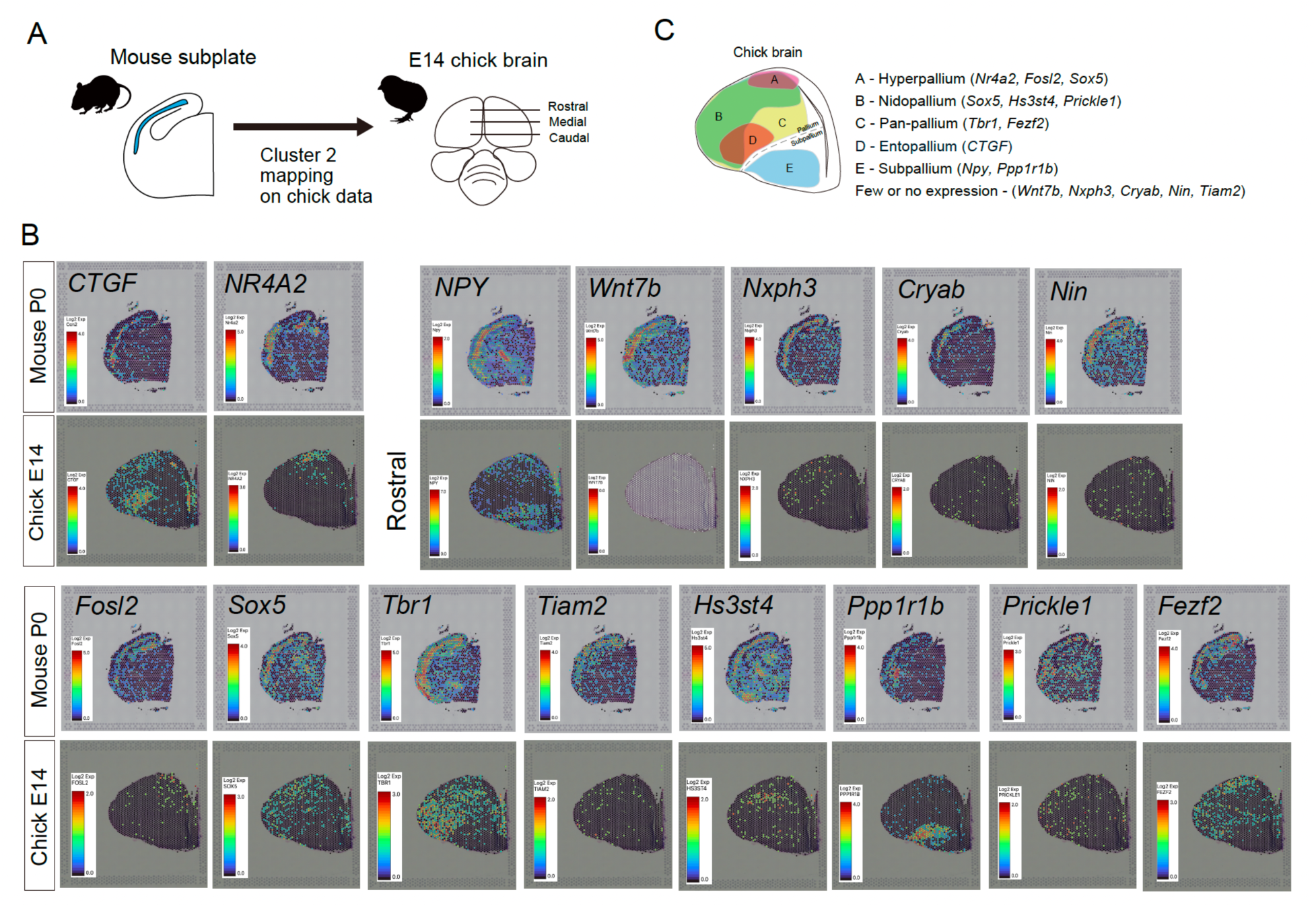
Expression patterns of mouse cluster 2 genes in mouse cortex and chick pallium. (**A**) Schematic overview of the cross-species mapping strategy. Genes enriched in the mouse subplate/L6-associated cluster 2 were selected from the mouse Visium dataset and their expression patterns were examined in the chick E14 pallium using the chick Visium dataset. (**B**) Spatial expression patterns of representative mouse cluster 2 genes in the mouse P0 cortex and chick E14 pallium. For each gene, the upper panel shows expression in the mouse P0 Visium dataset, and the lower panel shows the corresponding expression pattern in the chick E14 Visium dataset. Chick sections shown here correspond to the rostral level. (**C**) Summary of spatial expression domains of mouse cluster 2 genes in the chick E14 pallium. Genes were classified according to their predominant expression domains: hyperpallium, nidopallium, pan-pallial expression, entopallium, subpallium, or weak/no detectable expression.

**Fig. S8.**
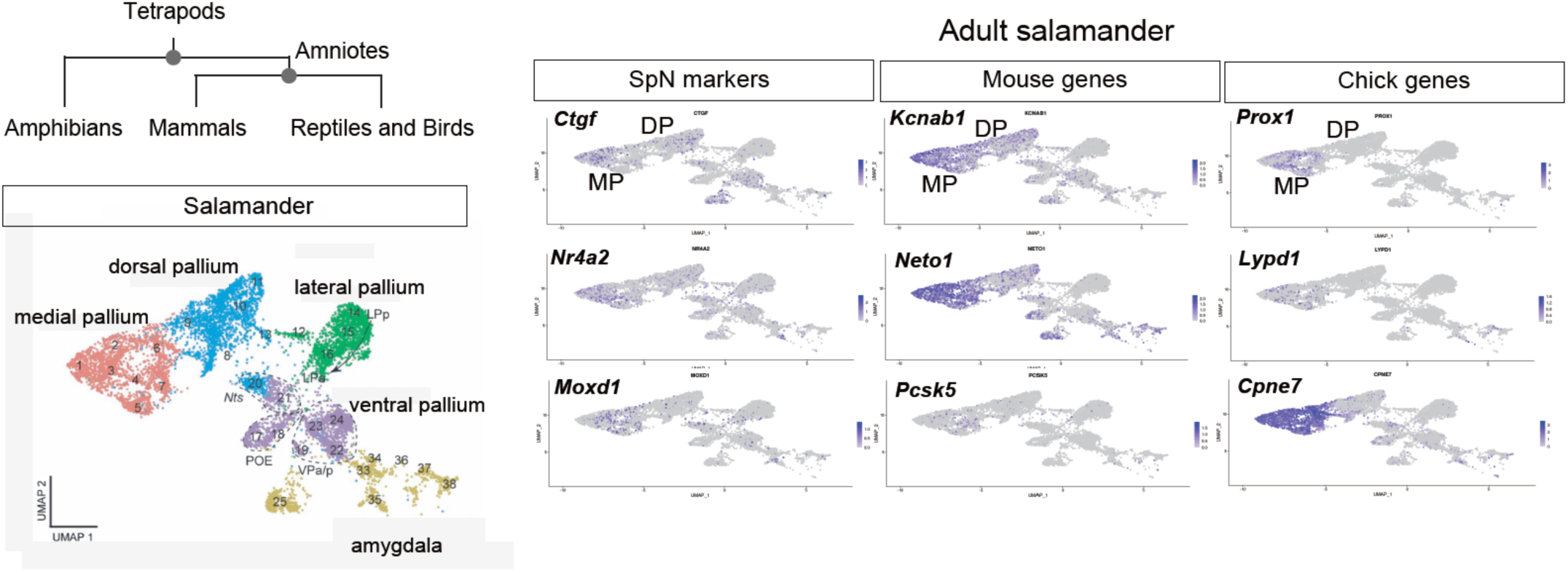
SpN marker expression in salamander and comparison with amniote signature. Left: Phylogenetic schematic indicating amphibians (salamander; *Pleurodeles waltl*) relative to amniotes. UMAP shows cell populations of the salamander pallium. Right: Feature plots showing expression of canonical SpN markers (e.g., Ctgf, Nr4a2, Moxd1) and representative mouse- and chick-associated marker genes in the adult salamander dataset. The UMAP in the left panel was retrieved and modified from Fig. 2A in *Woych et al. (2022)*, and the feature plots in the right panel were created by the web app provided by *Woych et al. (2022)* (https://toscheslab.shinyapps.io/salamander_telencephalon/).

**Fig. S9.**
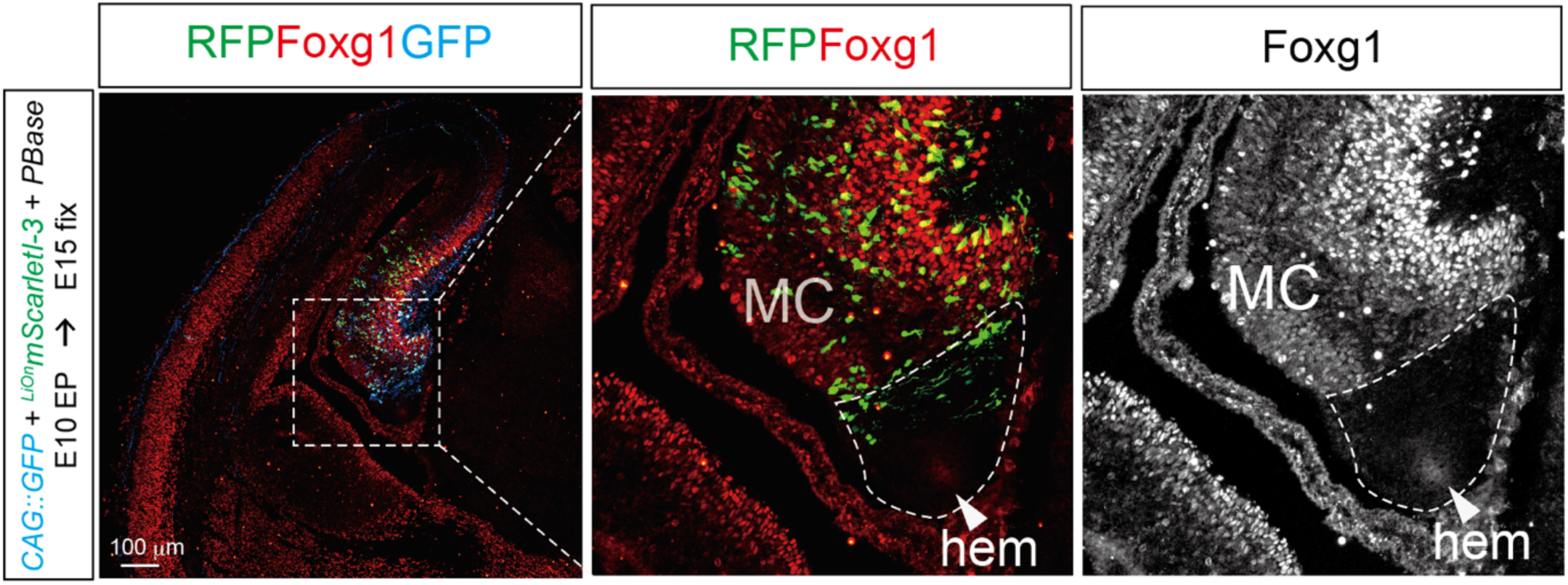
Plasmids electroporate both the Foxg1-positive medial cortex and the Foxg1-negative cortical hem. Representative coronal section of the embryonic mouse telencephalon electroporated at E10 with *CAG::GFP*, *^LiOn^CAG∞mScarletI-3*, and PBase, and analyzed at E15. Left: merged image showing GFP (blue), RFP (*^LiOn^CAG∞mScarletI-3*: green), and Foxg1 (red). Middle: GFP and RFP channels with the cortical hem region outlined (dashed line). Right: Foxg1 immunostaining indicating the Foxg1-positive medial pallium (MP) and Foxg1-negative cortical hem (hem; dashed outline). Scale bar, 100 μm.

**Fig. S10.**
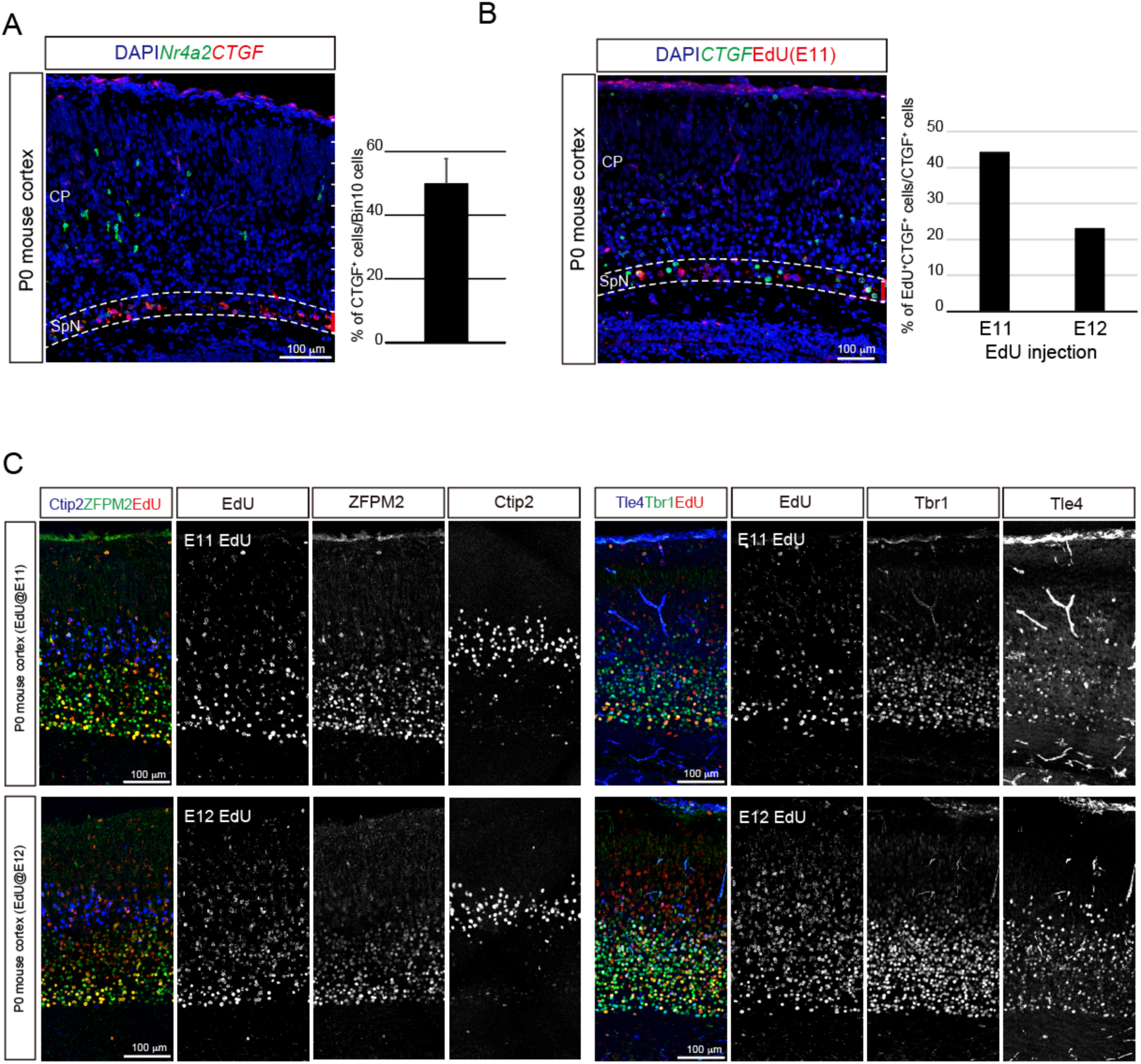
Birthdating analysis of early-born neurons in the mouse cortex. (**A**) Representative image of the P0 mouse cortex immunostained for Nr4a2 and CTGF (with DAPI). The dashed line delineates the subplate (SpN) region. Right: Quantification of the proportion of CTGF⁺ cells among Bin 10⁺ cells. (**B**) EdU birthdating of CTGF⁺ cells in the P0 mouse cortex. Pregnant mice were injected with EdU at E11 or E12, and brains were analyzed at P0 by CTGF immunostaining and EdU detection. The dashed line indicates the subplate neuron (SpN) region. Right: Percentage of EdU⁺CTGF⁺ cells among the total CTGF⁺ cells for each EdU time point. (**C**) Cell-type characterization of early-born (EdU-labeled) cortical neurons. Pregnant mice were injected with EdU at E11 or E12, and P0 cortices were stained for EdU together with neuronal subtype markers. Representative sections show EdU labeling with Ctip2 and ZFPM2 (left), and with Tbr1 and Tle4 (right), to assess the identities of early-born neurons. Scale bars, 100 μm.

**Fig. S11.**
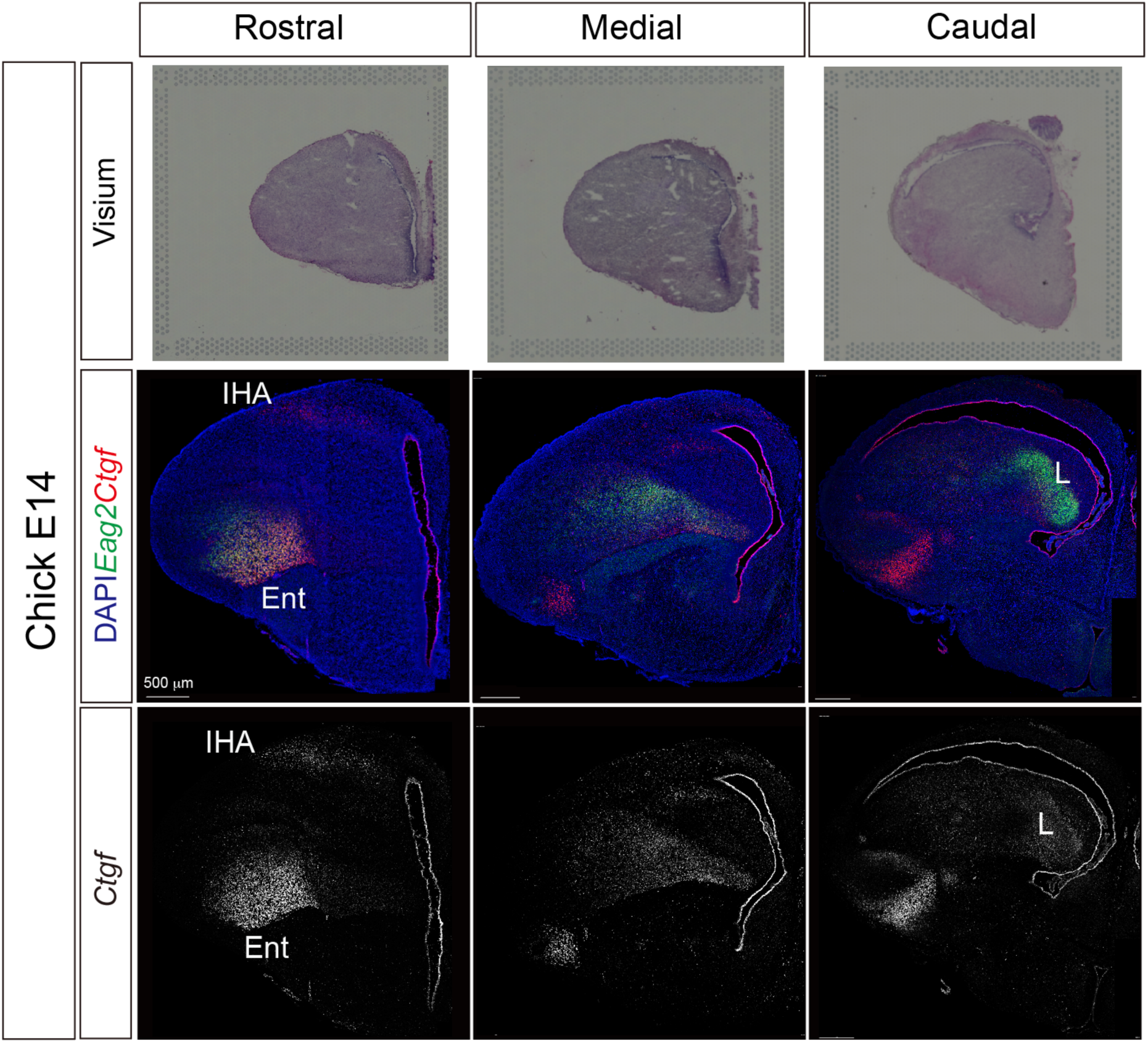
Rostral–caudal distribution of *Ctgf* expression in the chick E14 pallium. Representative coronal sections of the chick telencephalon at E14, shown at three rostral–caudal levels (rostral, medial, and caudal). Top row: Brightfield images. Middle row: Merged immunofluorescence images showing DAPI (blue), *Eag2* (green), and *Ctgf* (red) with the pallial region outlined. Bottom row: *Ctgf* channel shown alone to highlight the spatial distribution of the *Ctgf* signal across levels. ENT, entopallium; IHA, intermediate hyperpallium apicale; L, field L.

**Fig. S12.**
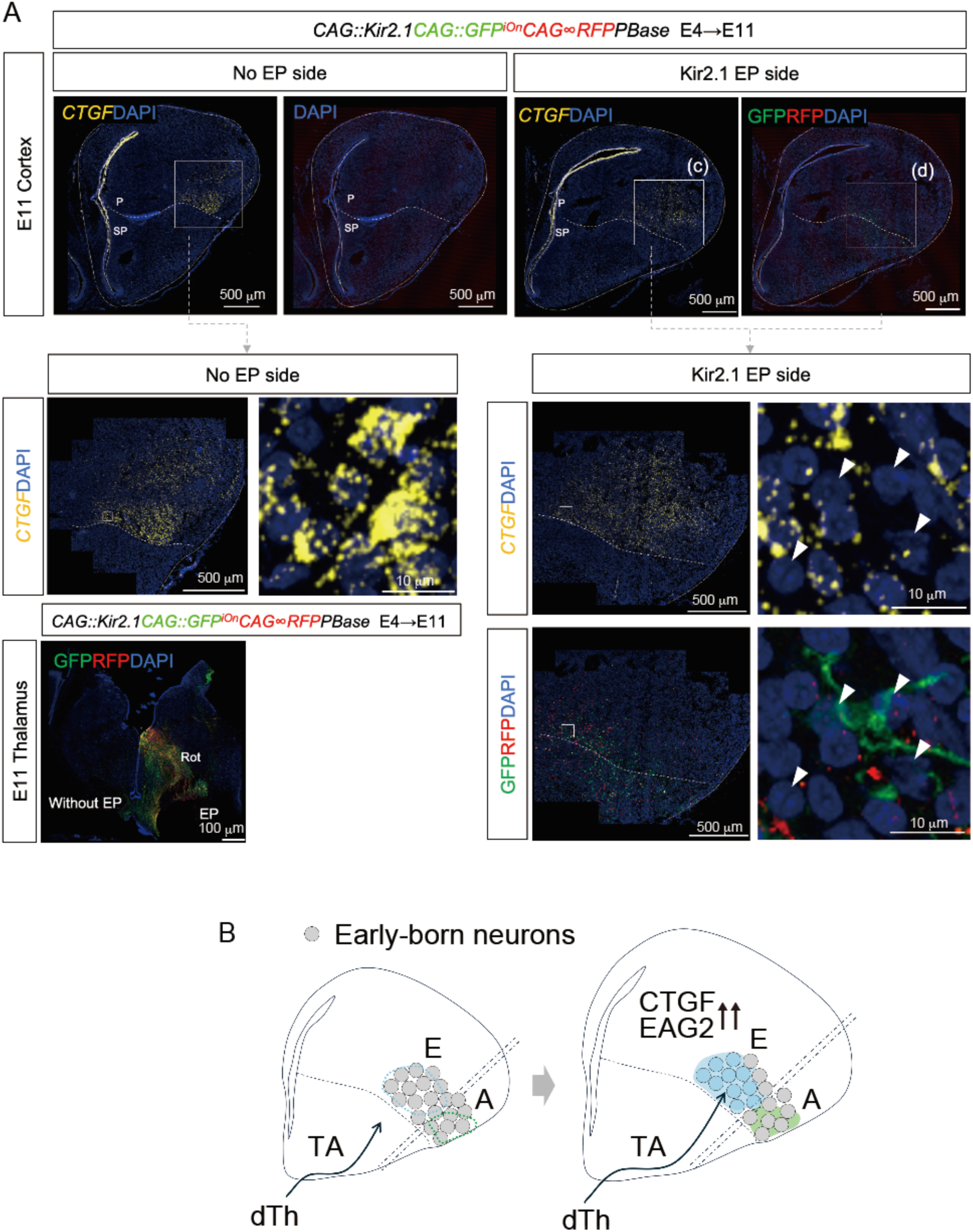
Thalamic input silencing by Kir2.1 alters Ctgf expression in the chick pallium. (**A**) Representative E11 chick telencephalon electroporated at E4 with *CAG::Kir2.1*, *CAG::GFP*, *^iOn^CAG∞RFP*, and *PBase*. Coronal sections show the non-electroporated (No EP) side and the Kir2.1-electroporated (EP) side. *CTGF* mRNA was detected by RNAscope (yellow) with DAPI counterstaining (blue), and GFP/RFP signals are shown as indicated. Boxed regions are shown at higher magnification. Arrowheads indicate cells located adjacent to GFP-positive thalamic axons in the electroporated condition. Bottom left: Thalamus images from the same brain showing GFP/RFP labeling on the EP side and the contralateral side; Rot, nucleus rotundus. (**B**) Schematic model illustrating that thalamic axonal input from the dTh modulates CTGF expression in early-born neuronal populations within the pallial target region. TA, thalamic axons; dTh, dorsal thalamus. Nuclei are counterstained with DAPI in all images. Scale bars, as indicated.

**Fig. S13.**
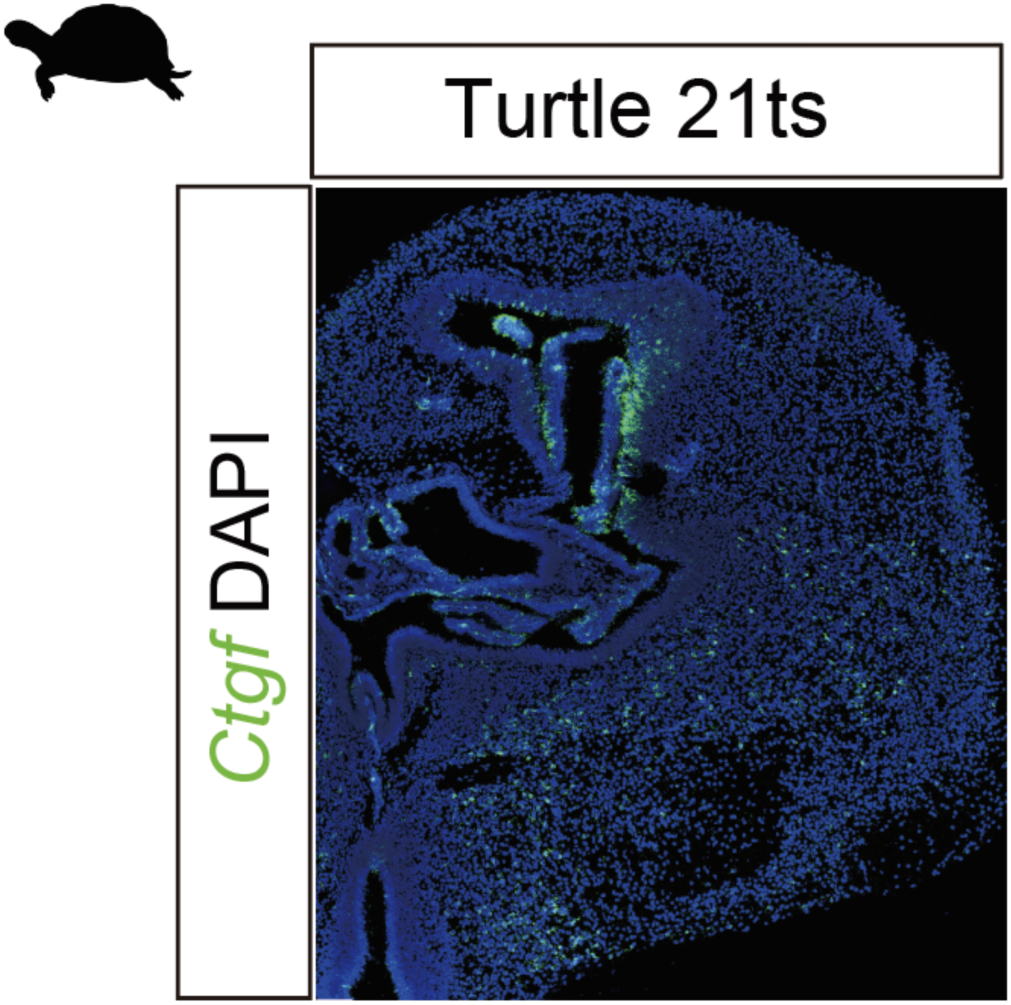
Rostral–caudal distribution of Ctgf mRNA in the turtle pallium at 21 ts. Representative coronal sections of the turtle telencephalon at stage 21 ts showing Ctgf mRNA detected by RNAscope (green) with DAPI counterstaining (blue). The caudal level is shown on the left, and the rostral level is on the right.

**Fig. S14.**
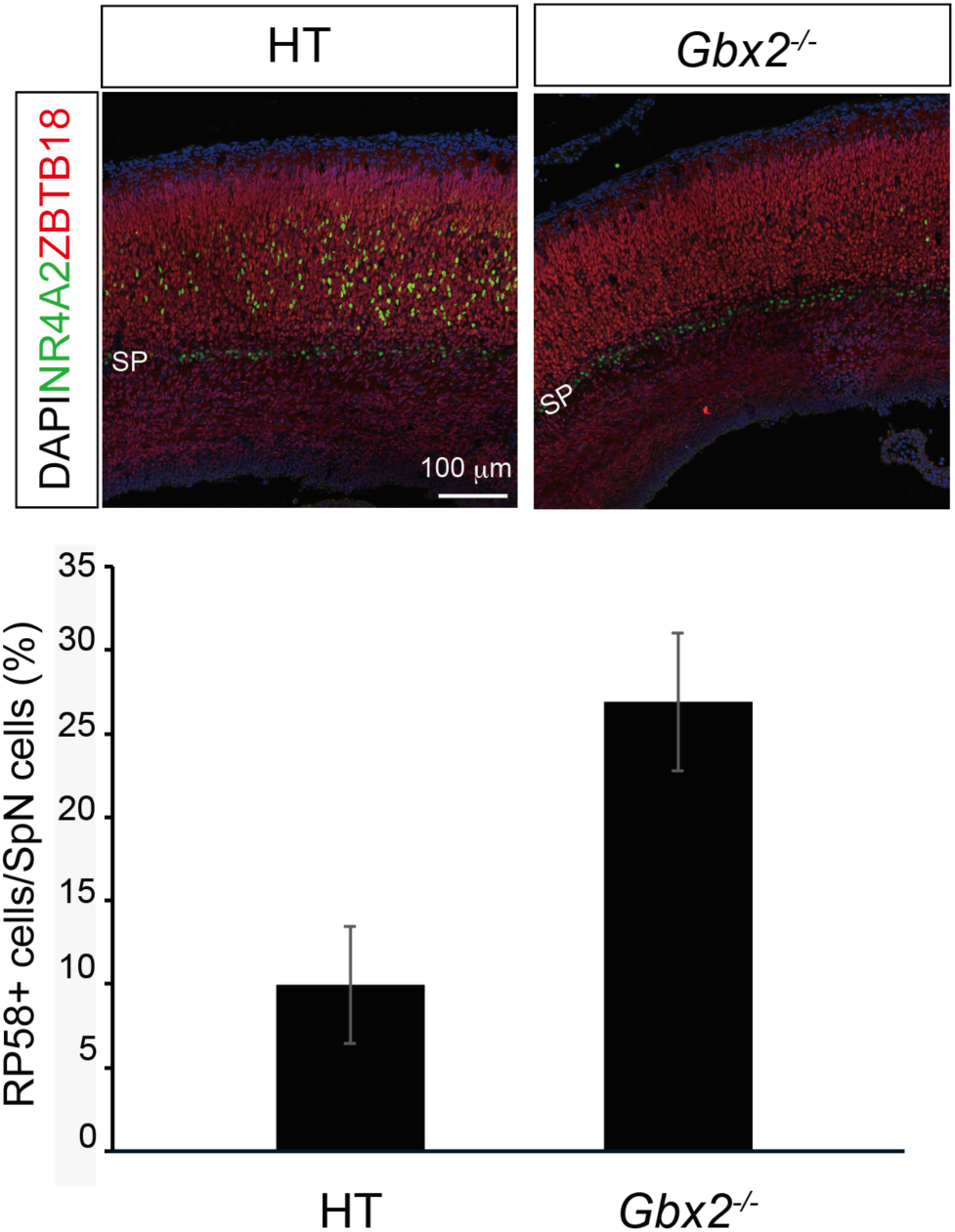
Increased ZBTB18 expression among subplate neurons in Gbx2 knockout cortex. Representative immunofluorescence images of cortical sections from heterozygous (HT) and *Gbx2*^−/−^ embryos stained for NR4A2 (green), ZBTB18 (red), and DAPI (blue). The subplate (SP) is indicated. Bottom: Quantification of the ZBTB18-positive cell fraction among subplate neurons (SpNs), showing an increased ZBTB18-positive proportion in *Gbx2*^−/−^ compared with HT. Scale bar, 100 μm.

**Fig. S15.**
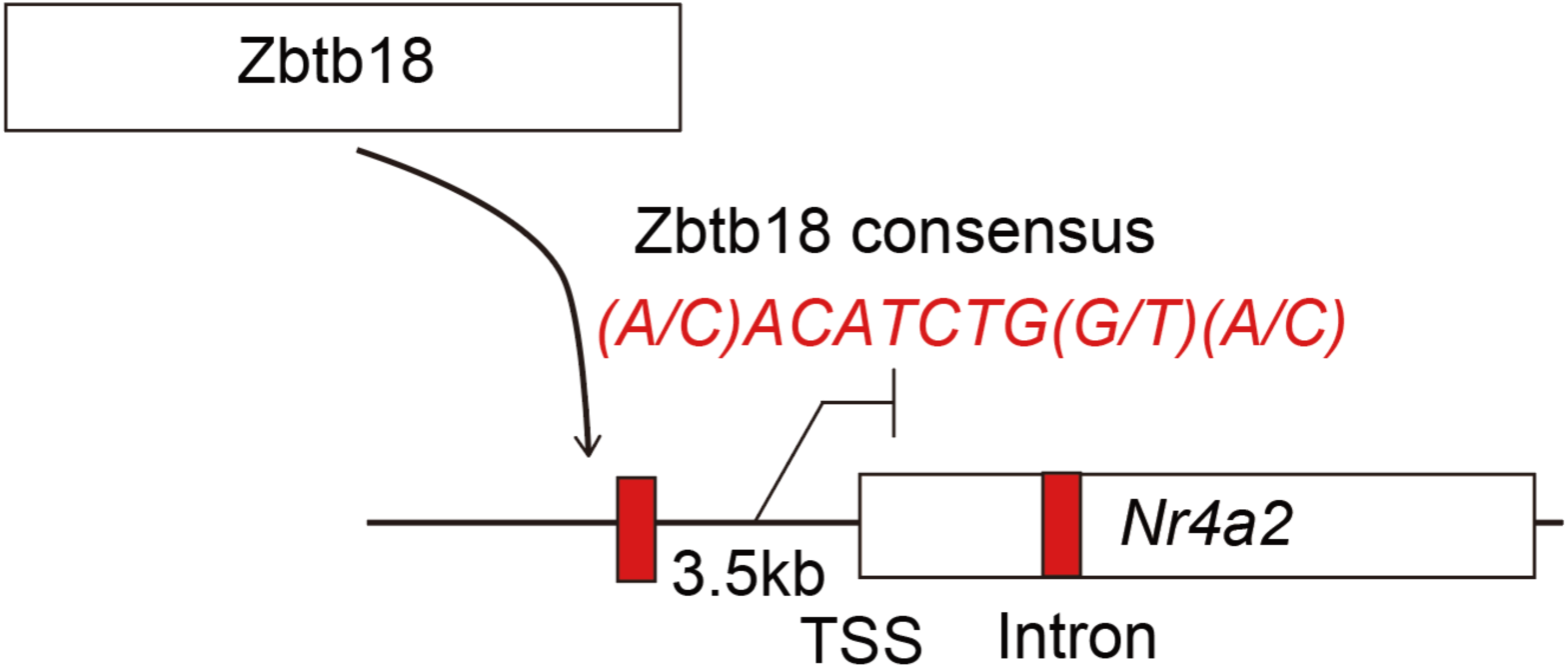
A putative Zbtb18-binding motif in the Nr4a2 locus. Schematic of the Nr4a2 genomic region showing a putative Zbtb18-binding site (red) located ∼3.5 kb upstream of the transcription start site (TSS). The Zbtb18 consensus motif is indicated in red. Boxes denote the promoter/TSS region and an intronic site within Nr4a2.

**Fig. S16.**
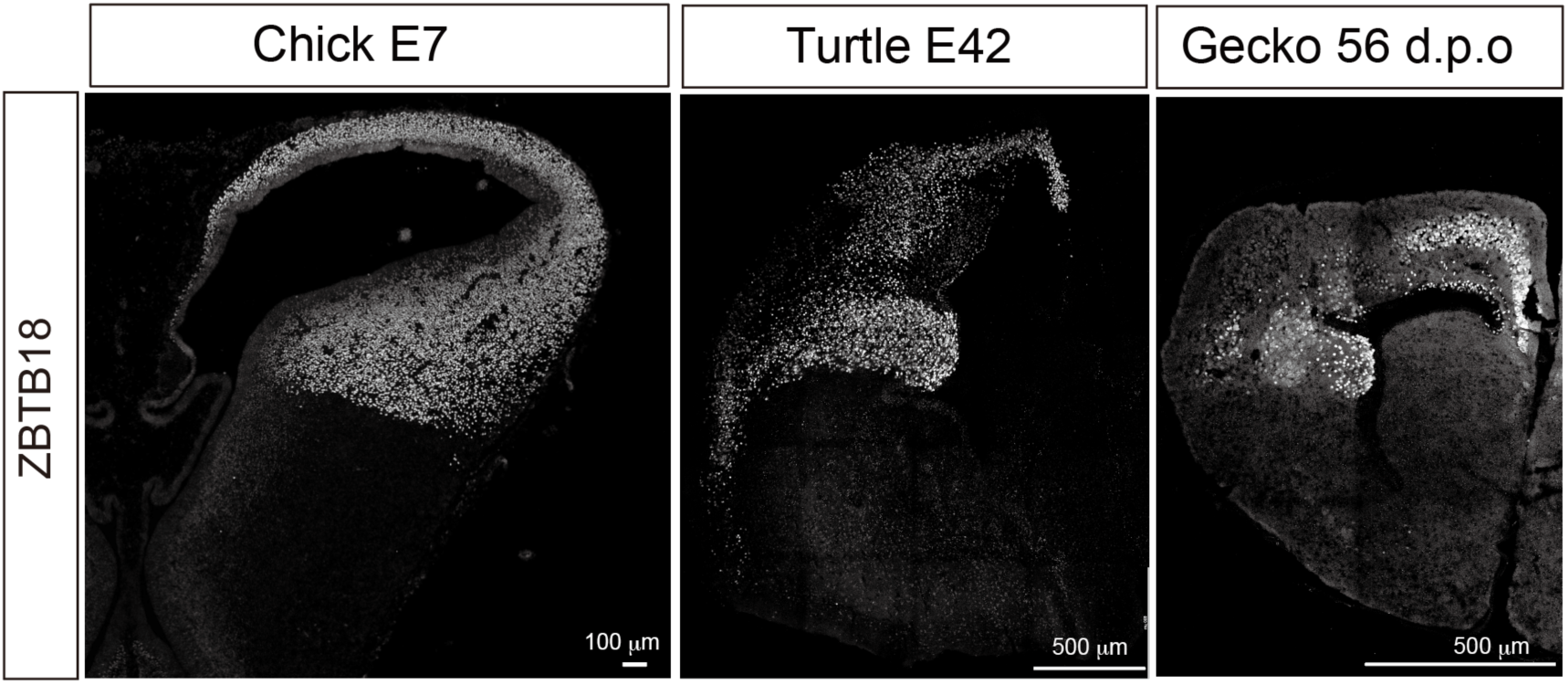
Conserved ZBTB18 expression in the developing pallium across sauropsids. Representative coronal sections showing Zbtb18 immunoreactivity in the pallia of chicks (E7), turtles (E42), and geckos (56 d.p.o.). Nuclei are counterstained with DAPI. Scale bars, as indicated.

## References and Notes

1. K. L. Allendoerfer, C. J. Shatz, The subplate, a transient neocortical structure: its role in the development of connections between thalamus and cortex. Annu Rev Neurosci 17, 185–218 (1994).

2. H. J. Luhmann, W. Kilb, I. L. Hanganu-Opatz, Subplate cells: amplifiers of neuronal activity in the developing cerebral cortex. Front Neuroanat 3, 19 (2009).

3. Z. Molnar, K. Y. Kwan, Development and Evolution of Thalamocortical Connectivity. Cold Spring Harb Perspect Biol 16, (2024).

4. Z. Molnar, H. J. Luhmann, P. O. Kanold, Transient cortical circuits match spontaneous and sensory-driven activity during development. Science 370, (2020).

5. C. Ohtaka-Maruyama, Subplate Neurons as an Organizer of Mammalian Neocortical Development. Front Neuroanat 14, 8 (2020).

6. C. Ohtaka-Maruyama et al., Synaptic transmission from subplate neurons controls radial migration of neocortical neurons. Science 360, 313–317 (2018).

7. A. Ghosh, C. J. Shatz, Involvement of subplate neurons in the formation of ocular dominance columns. Science 255, 1441–1443 (1992).

8. A. Hoerder-Suabedissen, Z. Molnar, Development, evolution and pathology of neocortical subplate neurons. Nat Rev Neurosci 16, 133–146 (2015).

9. P. O. Kanold, P. Kara, R. C. Reid, C. J. Shatz, Role of subplate neurons in functional maturation of visual cortical columns. Science 301, 521–525 (2003).

10. P. Rakic, Evolution of the neocortex: a perspective from developmental biology. Nat Rev Neurosci 10, 724–735 (2009).

11. R. L. Reep, Cortical layer VII and persistent subplate cells in mammalian brains. Brain Behav Evol 56, 212–234 (2000).

12. T. Fujita et al., The chick pallium displays divergent expression patterns of chick orthologues of mammalian neocortical deep layer-specific genes. Sci Rep 9, 20400 (2019).

13. J. F. Montiel et al., Hypothesis on the dual origin of the Mammalian subplate. Front Neuroanat 5, 25 (2011).

14. W. Z. Wang et al., Comparative aspects of subplate zone studied with gene expression in sauropsids and mammals. Cereb Cortex 21, 2187–2203 (2011).

15. Y. Hara et al., The spatial transcriptome of the late-stage embryonic and postnatal mouse brain reveals spatiotemporal molecular markers. Sci Rep 15, 12299 (2025).

16. J. Woych et al., Cell-type profiling in salamanders identifies innovations in vertebrate forebrain evolution. Science 377, eabp9186 (2022).

17. T. Kumamoto et al., Direct Readout of Neural Stem Cell Transgenesis with an Integration-Coupled Gene Expression Switch. Neuron 107, 617–630 e616 (2020).

18. M. Pedraza, A. Hoerder-Suabedissen, M. A. Albert-Maestro, Z. Molnar, J. A. De Carlos, Extracortical origin of some murine subplate cell populations. Proc Natl Acad Sci U S A 111, 8613–8618 (2014).

19. P. O. Kanold, H. J. Luhmann, The subplate and early cortical circuits. Annu Rev Neurosci 33, 23–48 (2010).

20. R. Katayama, T. Kumamoto, K. Wada, C. Hanashima, C. Ohtaka-Maruyama, Thalamic activity-dependent specification of sensory input neurons in the developing chick entopallium. J Comp Neurol 532, e25627 (2024).

21. E. Rueda-Alana et al., Evolutionary convergence of sensory circuits in the pallium of amniotes. Science 387, eadp3411 (2025).

22. H. M. Tsai, B. B. Garber, L. M. Larramendi, 3H-thymidine autoradiographic analysis of telencephalic histogenesis in the chick embryo: II. Dynamics of neuronal migration, displacement, and aggregation. J Comp Neurol 198, 293–306 (1981).

23. S. Yoshinaga et al., Comprehensive characterization of migration profiles of murine cerebral cortical neurons during development using FlashTag labeling. iScience 24, 102277 (2021).

24. F. Aboitiz, J. Montiel, R. R. García, Ancestry of the mammalian preplate and its derivatives: evolutionary relicts or embryonic adaptations? Reviews in the Neurosciences 16, 359–376 (2005).

25. T. Kumamoto et al., Foxg1 coordinates the switch from nonradially to radially migrating glutamatergic subtypes in the neocortex through spatiotemporal repression. Cell Rep 3, 931–945 (2013).

26. I. Kostovic, The enigmatic fetal subplate compartment forms an early tangential cortical nexus and provides the framework for construction of cortical connectivity. Prog Neurobiol 194, 101883 (2020).

27. Z. Li et al., Adaptive evolution of gene regulatory networks in mammalian neocortex. Nature, (2026).

28. N. Kaneko et al., ADAMTS2 promotes radial migration by activating TGF-beta signaling in the developing neocortex. EMBO Rep 25, 3090–3115 (2024).

29. D. Arendt et al., The origin and evolution of cell types. Nat Rev Genet 17, 744–757 (2016).

## References

1. M. Ohtsuka, M. Sato, H. Miura, S. Takabayashi, M. Matsuyama, T. Koyano, N. Arifin, S. Nakamura, K. Wada, C. B. Gurumurthy, i-GONAD: a robust method for in situ germline genome engineering using CRISPR nucleases. Genome Biol. 19, 25 (2018).

2. K. Wada, R. Katayama, I. Aota, C. Hanashima, T. Kumamoto, C. Ohtaka-Maruyama, Lineage analysis of pallial subdivision-derived cells in the developing chick pallium. Dev. Growth Differ. 67, 466–478 (2025).

3. T. Kumamoto, F. Maurinot, R. Barry-Martinet, C. Vaslin, S. Vandormael-Pournin, M. Le, M. Lerat, D. Niculescu, M. Cohen-Tannoudji, A. Rebsam, K. Loulier, S. Nedelec, S. Tozer, J. Livet, Direct Readout of Neural Stem Cell Transgenesis with an Integration-Coupled Gene Expression Switch. Neuron 107, 617–630.e6 (2020).

4. A. C. Adey, Integration of Single-Cell Genomics Datasets. Cell 177, 1677–1679 (2019).

5. Y. Hara, T. Kumamoto, N. Yoshizawa-Sugata, K. Hirai, X. Song, H. Kawaji, C. Ohtaka-Maruyama, The spatial transcriptome of the late-stage embryonic and postnatal mouse brain reveals spatiotemporal molecular markers. Sci. Rep. 15 (2025).

6. A. Dobin, C. A. Davis, F. Schlesinger, J. Drenkow, C. Zaleski, S. Jha, P. Batut, M. Chaisson, T. R. Gingeras, STAR: ultrafast universal RNA-seq aligner. Bioinformatics 29, 15–21 (2013).

7. M. Pertea, G. M. Pertea, C. M. Antonescu, T.-C. Chang, J. T. Mendell, S. L. Salzberg, StringTie enables improved reconstruction of a transcriptome from RNA-seq reads. Nat. Biotechnol. 33, 290–295 (2015).

8. G. X. Y. Zheng, J. M. Terry, P. Belgrader, P. Ryvkin, Z. W. Bent, R. Wilson, S. B. Ziraldo, T. D. Wheeler, G. P. McDermott, J. Zhu, M. T. Gregory, J. Shuga, L. Montesclaros, J. G. Underwood, D. A. Masquelier, S. Y. Nishimura, M. Schnall-Levin, P. W. Wyatt, C. M. Hindson, R. Bharadwaj, A. Wong, K. D. Ness, L. W. Beppu, H. J. Deeg, C. McFarland, K. R. Loeb, W. J. Valente, N. G. Ericson, E. A. Stevens, J. P. Radich, T. S. Mikkelsen, B. J. Hindson, J. H. Bielas, Massively parallel digital transcriptional profiling of single cells. Nat. Commun. 8, 14049 (2017).

9. D. J. Di Bella, E. Habibi, R. R. Stickels, G. Scalia, J. Brown, P. Yadollahpour, S. M. Yang, C. Abbate, T. Biancalani, E. Z. Macosko, F. Chen, A. Regev, P. Arlotta, Molecular logic of cellular diversification in the mouse cerebral cortex. Nature 595, 554–559 (2021).

10. Y. Hao, S. Hao, E. Andersen-Nissen, W. M. Mauck 3rd, S. Zheng, A. Butler, M. J. Lee, A. J. Wilk, C. Darby, M. Zager, P. Hoffman, M. Stoeckius, E. Papalexi, E. P. Mimitou, J. Jain, A. Srivastava, T. Stuart, L. M. Fleming, B. Yeung, A. J. Rogers, J. M. McElrath, C. A. Blish, R. Gottardo, P. Smibert, R. Satija, Integrated analysis of multimodal single-cell data. Cell 184, 3573–3587.e29 (2021).

11. C. Hafemeister, R. Satija, Normalization and variance stabilization of single-cell RNA-seq data using regularized negative binomial regression. Genome Biol. 20, 296 (2019).

12. T. Stuart, A. Butler, P. Hoffman, C. Hafemeister, E. Papalexi, W. M. Mauck 3rd, Y. Hao, M. Stoeckius, P. Smibert, R. Satija, Comprehensive Integration of Single-Cell Data. Cell 177, 1888–1902.e21 (2019).

